# Target RNA-triggered CRISPR-Cas12a2 Preferentially Cleaves Collateral DNA over RNA

**DOI:** 10.1101/2025.01.05.631166

**Authors:** Sobita Kunwar, Thomson Hallmark, Sudeshna Manna, Dylan Keiser, Bronson Naegle, Aaron Thomas, Chase L. Beisel, Ryan N. Jackson

## Abstract

CRISPR-Cas systems often rely on collateral cleavage of nucleic-acid substrates to combat recognized mobile genetic elements. Of the CRISPR-associated (Cas) RNA-guided effector nucleases, Cas12a2 stands out as the only known example exhibiting rapid collateral cleavage of three distinct substrates: single-stranded (ss)RNA, ssDNA, and double-stranded (ds)DNA, after activating upon binding cognate RNA. However, little is known about the underlying mechanisms of collateral cleavage. Here, we show, using enzyme kinetics and inhibition assays, that Cas12a2 preferentially cleaves collateral DNA over RNA substrates, even when RNA substrates are more abundant. Additionally, using enzyme mutants, enzyme kinetics, and plasmid cleavage assays, we determine that the dsDNA cleavage mechanism relies on the ‘aromatic clamp’ residues that stabilize unwound and distorted dsDNA in the RuvC nuclease active site. Leveraging the cleavage preference for collateral DNA, we demonstrate that RNA-activated Cas12a2 can readily cleave a ssDNA probe in the presence of high concentrations of non-target RNA, while an RNA-targeting Cas13a cannot. This work provides foundational kinetic and biochemical insights into the collateral cleavage mechanism and substrate preferences of Cas12a2, with immediate implications for understanding Cas12a2-based immunity and developing Cas12a2-based technologies.

## Introduction

CRISPR-Cas12a2 is a single-subunit, type V CRISPR nuclease that defends bacteria against foreign nucleic acids (like viruses and plasmids) by targeting RNAs that are complementary to a bound CRISPR RNA (crRNA) and contain a 3′-end Protospacer Flanking Sequence (PFS) (1). RNA targeting induces a large conformational change in Cas12a2 that opens the RuvC nuclease domain (2). Once open, the nuclease non-specifically cleaves collateral ssRNA, ssDNA, and dsDNA, causing growth arrest of the host cell and stalling replication of the foreign nucleic acid (1). While other single-subunit Cas nucleases can collaterally cleave ssDNA or RNA (3–5), and some Cas12a1 enzymes have demonstrated the ability to cleave short dsDNA probes with free ends (6–11) and longer dsDNA sequences with ssDNA looped structures or 3′-end overhangs (10, 12, 13), Cas12a2 is the only known single-subunit CRISPR nuclease that rapidly degrades longer lengths of ssRNA, ssDNA and dsDNA, including supercoiled plasmids (1, 14). Furthermore, the Cas12a2-mediated cleavage of collateral dsDNA has been proposed as the major driver of the growth arrest phenotype correlated with Cas12a2-activated immunity (1, 2), while the indiscriminate collateral DNase activity of Cas12a1 has been shown to be non-essential for immunity (15).

While collateral dsDNA cleavage is hypothesized to drive growth arrest, the substrate preference of Cas12a2 for collateral DNA or RNA substrates is unclear, and it is unknown how Cas12a2 collateral nuclease activities are influenced by the presence of competing collateral substrate types. In particular, little is known about how collateral cleavage activities are influenced in a cellular environment with all three collateral substrate types available. Because the collateral nuclease activities of Cas12a2 suggest unique biological mechanisms and novel applications in which both RNA and DNA would be present (e.g., RNA diagnostics using DNA probes), we aimed to better understand the Cas12a2 collateral cleavage mechanisms in isolation and in the presence of multiple collateral substrate types.

Using steady-state kinetics, inhibition assays, and endpoint fluorescence assays, we determined that Cas12a2 shows a kinetic preference for cleaving collateral DNA substrates, even in the presence of excess RNA. Additionally, we show that the aromatic clamp residues that stabilize melted and bent dsDNA in the RuvC nuclease active site (2) play important but distinct roles in dsDNA cleavage. Leveraging the cognate RNA-triggered preference for cleaving collateral DNA, we then demonstrate that when combined with a DNA probe, RNA-guided Cas12a2 can detect specific RNA targets in a sample with high concentrations of non-target RNA without target pre-amplification or reverse transcription, consistent with work reported while this manuscript was under revision (14, 16). Collectively, this work reveals kinetic mechanisms unique to Cas12a2, provides insight into how Cas12a2 drives growth arrest during an immune response, and deepens understanding of Cas12a2 collateral cleavage mechanisms.

## Methods

### Substrates and plasmids

All nucleotide substrates described in **Supplementary Table 1** and the LbCas12a1enzyme were purchased from Integrated DNA Technologies (IDT). Plasmid constructs used for expression and purification of Cas12a2 and Cas12a2 mutants were the same as those previously described (1, 2).

### Expression and initial purification of SuCas12a2

Cas12a2 from *Sulfuricurvum* sp. PC08-66 (SuCas12a2) and the mutants thereof were expressed and purified as described previously (1, 2). Briefly, Cas12a2 and mutant expression plasmids were freshly transformed into BL21 NiCo (DE3) cells, and a single colony was used to grow 20 mL overnight cultures. Overnight cultures were used to inoculate 1.0 L of TB media. Cells were grown to an OD600 of 0.6-0.8 and cold-shocked for 15 minutes on ice before induction with 0.1 M IPTG. The temperature was lowered to 18°C, and cells were grown for 16-18 hours. Cells were harvested by centrifugation at 3000 rpm for 30 min.

Cell pellets were resuspended in lysis buffer (25 mM Tris pH 7.2, 500 mM NaCl, 2 mM MgCl_2_, 10 mM imidazole, 10% glycerol), treated with protease inhibitors (0.5µg/mL aprotinin, 0.5µg/mL leupeptin, 1mM AEBSF (4-(2-Aminoethyl)benzenesulfonyl fluoride hydrochloride), and 0.7µg/mL pepstatin A) and 1 mg/mL lysozyme, and allowed to incubate on ice for 30 minutes. Cells were lysed by sonication and clarified by centrifugation at 15000 rpm for 25 min at 4°C. The lysate was batch-bound to 2 mL of nickel resin for 30 min at 4 °C, then flowed through the resin three times. Resin was washed with 500 mL of nickel wash buffer (25 mM Tris pH 7.2, 2 M NaCl, 2 mM MgCl_2_, 10 mM imidazole, 10% glycerol), then eluted in 4 mL fractions with elution buffer (25 mM Tris pH 7.2, 500 mM NaCl, 2 mM MgCl_2_, 500 mM imidazole, 10% glycerol).

Nickel elutions were transferred to low salt buffer (25 mM Tris pH 7.2, 50 mM NaCl, 2 mM MgCl2, 10% glycerol) using a HiPrep 26/10 desalting column (Cytiva), then loaded over a 5 mL HiTrap SP HP column (Cytiva), washed with 10% high salt buffer (25 mM Tris pH 7.2, 1.0 M NaCl, 2 mM MgCl_2_, 10% glycerol) and eluted with gradient elution to 100% high salt buffer over 5 column volumes. Peak fractions were pooled and concentrated with 100 kDa MWCO spin concentrators (Corning) to a volume of about 1 mL and then run over a Size Exclusion Column (SEC), for example, Superdex 200 10/300 increase GL sizing column (Cytiva), equilibrated with SEC Buffer (12.5LJmM HEPES pHLJ7.2, 150 mM NaCl, 2LJmM MgCl_2_). Peak fractions were pooled and concentrated using 100 kDa MWCO spin concentrators (Corning), aliquoted, flash-frozen in liquid nitrogen, and stored at −80°C.

### Preparation of nucleic acid substrates

Oligos (**Supplementary Table 1**) were resuspended to 1.0 mM in water. DNA oligos were stored at −20°C while RNA samples were aliquoted and stored at −80°C. All oligos were quantified before each use by spectrophotometry using an ND-1000 spectrophotometer (NanoDrop), measured at ng/µL, and converted to µM using an online “Weight to Molar Quantity (for nucleic acids)” tool (www.bioline).

### Complex Formation

Binary complex was formed by combining protein and synthetic crRNA in a 1:1.2 molar ratio in SEC buffer (25mM HEPES, pH 7.2, 150 mM KCl, 2 mM MgCl_2_) and incubating at 24°C for 20 minutes (2). The binary complex was then further purified over a Superdex 200 10/300 increase GL sizing column (Cytiva) into SEC Buffer (12.5LJmM HEPES pHLJ7.2, 150 mM NaCl, 2LJmM MgCl_2_), to remove excess crRNA. The eluted protein complex was concentrated in a 100-kDa MWCO spin concentrator (Corning) aliquoted, flash frozen in liquid nitrogen, and stored at - 80°C.

The ternary complex was assembled by first incubating the protein with crRNA at a 1:1.2 molar ratio in SEC buffer (25 mM HEPES, pH 7.2; 150 mM KCl; 2 mM MgClLJ) at 24°C for 20 minutes. Target RNA was then added in excess of protein and crRNA at a ratio of 1:1.2:1.5 (protein:crRNA:target RNA), followed by a second incubation at 24 °C for 20 min to allow ternary complex formation. To remove excess crRNA and target RNA, the ternary complex was purified over a Superdex 200 10/300 increase GL sizing column (Cytiva) into SEC Buffer (12.5LJmM HEPES pHLJ7.2, 150 mM NaCl, 2LJmM MgCl_2_) **(Supplemental Figure S1)**. Eluted ternary complex was concentrated in a 100-kDa MWCO spin concentrator (Corning), aliquoted, flash frozen in liquid nitrogen, and stored at −80°C.

### Fluorescence anisotropy binding assay

#### Target binding assay

Binary complex was two-fold serially diluted into a 1X binding buffer (25LJmM HEPES pHLJ7.2, 150mM KCl, 2LJmM MgCl_2_, 5μg/ml BSA) for 11 samples, with a 12th sample containing only binding buffer. Binary complex samples were combined with 5′-FAM-labeled phosphorothioated (non-hydrolyzable) target RNA probe in a black-bottom, 96-well plate (Corning) for a final concentration of 10 nM RNA probe. A 2-fold serial dilution series of Cas12a2 binary in a range of nanomolar concentrations (875, 437, 219, 109, 55, 27, 14, 7, 4, 2, 1, and 0.0 nM) in binding buffer. The delta anisotropy was measured in a Synergy H4 Hybrid plate reader (BioTek) at multiple time points to determine when Bmax and Kd stop fluctuating, indicating the time point at which the reaction reaches binding equilibrium (**Supplemental Figure S2**). All reported Kds were calculated using the delta anisotropy values measured at 30 minutes, as all substrates appeared to have reached binding equilibrium at that time. Kds were determined by fitting data to a quadratic-formula-based binding equation (see data collection and processing).

#### Non-target (Collateral) Binding Assay

Ternary complex was two-fold serially diluted into a binding buffer for 11 samples, with a 12th sample of only binding buffer (nanomolar concentrations of 875, 437, 219, 109, 55, 27, 14, 7, 4, 2, 1 and 0.0 nM) ternary complex in binding buffer. Ternary complex samples were then combined with a 5′-FAM-labeled non-target phosphorothioated (non-hydrolyzable) substrate probe in a black-bottom 96-well plate (Corning) at a final probe concentration of 10 nM. The plate was measured for delta anisotropy at different time points to ensure the binding reached equilibrium in a Synergy H4 Hybrid plate reader (BioTek) (**Supplemental Figure S2**). All reported Kds were calculated using the delta anisotropy values measured at 30 minutes, as all substrates appeared to have reached binding equilibrium at that time (**Supplemental Figure S2**). Kds were determined by fitting the data to a quadratic formula (see data collection and processing).

#### Data collection and processing

The plate reader was set to collect fluorescence emission end-point polarization with wavelengths λex. = 485/20 nm and λem. = 528/20 nm. The 20 nm in the denominator is the band gap filter used. The detector was set to 4.0 mm above the plate, and individual gain values were calculated for each substrate using the auto-gain function in Gen5 software (BioTek). The Gen5 software was used to output fluorescence polarization and calculate anisotropy values.

Three independent replicates were collected for each protein concentration, and the average anisotropy of three no protein complex conditions was calculated as the background signal. Background was subtracted from each data point to calculate the reported ΔAnisotropy. Binding isotherms were generated by fitting the data against the quadratic binding equation (17) in GraphPad Prism:

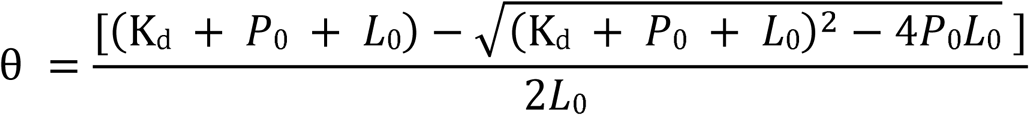

where θ is the fraction of nucleic acid substrate bound, K_d_ is the dissociation constant, L_0_ is the total nucleic acid substrate concentration (e.g., 10 nM), and P_0_ is the total protein complex concentration (apo, binary, or ternary) at each data point. The mean and standard error from three independent experiments are reported. As a control to compare substrate binding to another protein other than Cas12a2, a similar methodology was used to determine Kd values with Bovine Serum Albumin (**Supplemental Figure S3**). Also, a similar methodology was used to measure the binding of crRNA and FAM-labeled target RNA, as a control.

### Fluorescence reporter kinetic assay

Fluorescent reporter substrates (**Supplemental Table 1**) were diluted in 1x NEB3.1 buffer (50 mM Tris-HCl, pH 7.9, 100 mM NaCl, 10 mM MgCl_2_, 100 μg/mL bovine serum albumin (BSA)) to different concentrations ranging from 6µM to 0.1 nM, see **Supplemental Figure S4** for concentrations used in each assay. Reactions were initiated by adding 2x SEC-purified ternary complex to a final concentration of 15 nM. Reactions were combined in black bottomed 384-well plate (Greiner Bio-One).

Progress curves were collected using a Synergy H4 Hybrid plate reader, measuring fluorescence intensity at 19-second intervals using λex. = 485/20 nm and λem. = 528/20 nm with read height set to 8.0 mm and gain of 75. Fluorescence intensity was converted to concentration using standard curves for each substrate (see method for standard curve creation below and **Supplemental Figure S5**). Three independent replicates were collected for each condition, and slopes (steady-state velocities) of the linear portion of the progress curve (first six data points, each taken in a 19-second time interval) were determined by linear fit in GraphPad Prism. Kinetic parameters of Cas12a2 cleavage were determined by plotting the calculated steady state velocities (V_0_) versus the substrate concentration and fitting the data to the Michaelis-Menten equation in GraphPad Prism. The accuracy of the fit was calculated using confidence interval values (lower and upper limit) and was reported as a standard deviation for the V_max_, K_m_, k_cat_, and k_cat_/K_m_ values.

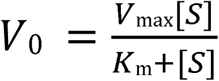

Catalytic efficiency (specificity constant) was determined to be the ratio of k_cat_/K_m_. The formula used to calculate k_cat_ is as follows, where [ES] is assumed to be equal to the total concentration of SEC-purified ternary Cas12a2 enzyme in the experiment at substrate-saturating conditions (i.e., concentrations that exhibit a rate of V_max_).

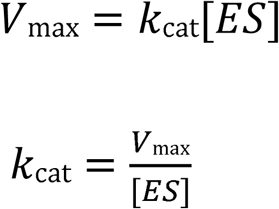

### Pairwise competition cleavage kinetics

Pairwise competition assays followed the same preparation of active ternary complex and reporter substrates described above. A 4x solution of phosphorothioated inhibitor was prepared in 1x NEB 3.1 buffer for each inhibitor concentration. Inhibitor concentrations were selected by screening several potential inhibitor conditions, resulting in final inhibitor concentrations ranging from 3 μM to no inhibitor. The active ternary complex, fluorescent reporter, and inhibitor were combined in a black-bottom 384-well plate at the same concentrations used for the kinetic assays. Reaction progress curves and data processing followed the protocol described above for the fluorescent reporter kinetic assays. Apparent kinetic values reported in **Table 3** were determined by fitting data to the Michaelis-Menten equation as described above.

Fitting different models of inhibition was performed in GraphPad Prism using the equation for competitive inhibition:

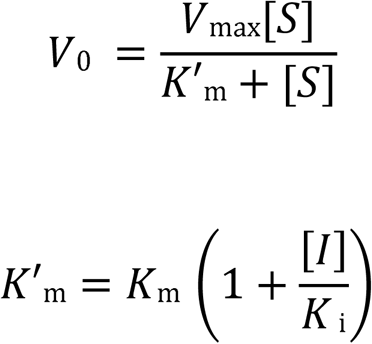

and mixed-model inhibition:

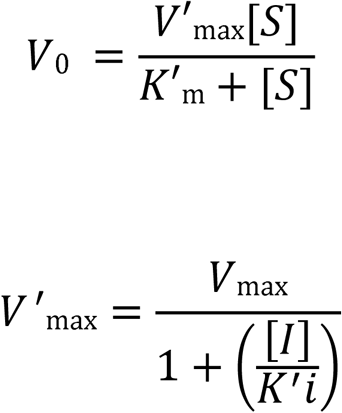

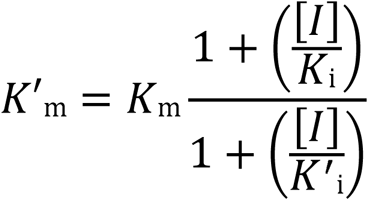

### Standard curve generation

Each fluorescent reporter substrate was combined with an active Cas12a2 ternary complex to a final concentration of 10 µM reporter and 15 nM ternary complex in 1x NEB 3.1 buffer. Samples were allowed to incubate overnight (16 - 18 hours) for 100% reporter cleavage and were then diluted into 1x NEB 3.1 buffer to form a standard curve (**Supplementary Figure S5**).

Fluorescence intensity was measured using a Synergy H4 Hybrid plate reader using λex. = 485/20 nm and λem. = 528/20 nm with read height set to 8.0 mm and gain of 75. A linear fit of the standard curves was prepared in GraphPad Prism (**Supplementary Table 3**), and the resulting fit-line was used to convert the observed fluorescence signal to a molar concentration in kinetic reporter assays using the following equation, where *I_fluor_* is the intensity of the fluorescence signal, and *Y_int._* and *Slope* are the y-intercept and slope of the best-fit line.

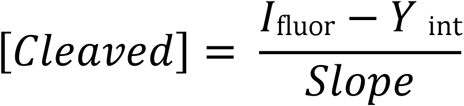

### Plasmid Cleavage Assay

A 100 μl reaction containing 14LnM ternary complex, 7LnM of pUC19 plasmid in 1× NEB 3.1 buffer was incubated at room temperature. At the indicated time points, 15 μl of the reaction was removed and quenched with phenol, and phenol–chloroform extraction was performed. The reactions were visualized on 1% agarose with ethidium bromide.

### Next-generation sequencing of plasmid cleavage

Plasmid cleavage reactions were prepared as described previously (1, 2). Briefly, 14 nM Cas12a2 was combined with 14 nM crRNA, 25 nM target RNA, and 7 nM supercoiled pUC19 plasmid in NEB 3.1 buffer (50 mM Tris-HCl, pH 7.9, 100 mM NaCl, 10 mM MgCl_2_, 100 μg/ml BSA). Reactions were run for 2 min before quenching in pH 8.0 phenol-chloroform. The aqueous layer from the phenol-chloroform extraction was combined with an equal volume of 50% glycerol and then separated on a 1% agarose gel. Bands in the agarose gel were visualized by EtBr on a UV-transilluminator (BioRad), and bands representing the linear species were extracted using a gel extraction kit (Synthbio).

The cleaved plasmids were first end-repaired using the NEB Ultra II End Repair/dA-Tailing Module. This polished the ends and added a 3′-A overhang to prevent ligation of fragments to each other and to aid adapter ligation. A biotinylated double-stranded i5 adapter was then ligated to the 5′ end. The sample was fragmented into 400-bp fragments using a Covaris S220 sonicator and cleaned up with magnetic beads. The end repair process was repeated, followed by ligation of a double-stranded i7 adapter to a 5′-phosphorylated i7-R. The DNA was then bound to streptavidin magnetic beads (Dynabeads MyOne Streptavidin C1), and unbound fragments were washed away. The DNA on the beads was then amplified using Q5 Hot Start and Illumina indexing primers. The libraries were then sequenced on the Illumina MiSeq using the 300-cycle v2 chemistry kit. Raw data sets were deposited at NCBI Sequence Read Archive (SRA) under submission SUB14940624 and are publicly available.

To generate a list of cut sites, reads were mapped to the pUC19 plasmid sequence. Then bedtools was used to create a table of the 5′ end of each mapped read. To account for potential errors from reads crossing the origin of the reference sequence, reads that cross the origin were separated and mapped to a second reference, and 5′ ends were identified with bedtools. Reads mapped to each location were then combined between the two sets of cut-sites, and read depth at each position was plotted in GraphPad Prism. To generate WebLogos of the cleavage consensus sequences, the 14-bp region around all 69 unique cut sites (6147 reads) was submitted to the WebLogo Server (UC Berkley)(18), and all reads were included. To eliminate sequence weighting from the most abundant site, we generated a second WebLogo without that site included.

### Determination of ssDNA reporter sequence with optimal signal-to-noise ratio

Active ternary complexes were prepared by incubating a mixture of Cas12a2 (1 µM), targeting or non-targeting crRNA (0.5 µM), and target RNA (50 nM) in 1 x NEB 3.1 buffer (50 mM Tris-HCl, pH 7.9, 100 mM NaCl, 10 mM MgCl_2_, 100 ug/mL BSA) for 30 min. All 6-mer ssDNA probes with 5′ FAM and 3′ 3IABkFQ of 10 µM stock concentration were added to the mixture to get a final concentration of 1 µM. The reaction mixtures were placed in a Polystyrene transparent conical base 96-well plate (SARSTEDT). Fluorescence intensity was measured at 29 °C using a BioTek Synergy H1 microplate reader with λ_ex._ = 484 nm and λ_em._ = 530 nm, read height 7 mm (bottom read), a gain value of 70 over a time period of 2 hours.

### Comparison of RNA inhibition on Cas12a2 and Cas13a

A 2x detection master mix was prepared with 2x NEB 3.1, 0.2 µM Nuclease, 0.24 µM crRNA, and 2 µM probe, where the nuclease is either SuCas12a2 or LwaCas13a (MCLabs), and the probe is either 6-nt poly T or poly U, respectively, labeled with 5′ fluorescein and 3′ IowaBlack quencher (IDT). A stock of 0.1 µM target RNA was prepared with a range of concentrations (2000, 200, 20, 2.0, 0.2, 0.02, and 0.0 ng/µl) of total yeast RNA (Fisher). For additional controls, the 2x detection master mix (5 µl) was combined with 5 µl of target RNA with or without total yeast RNA. Samples were incubated at 37°C for 30 minutes before being measured for endpoint fluorescence using a Synergy H4 Hybrid plate reader with λex. = 485/20 nm and λem. = 528/20 nm, read height set to 8.0 mm, and a gain of 65. The average fluorescence intensity of samples containing target RNA but no yeast RNA, for both SuCas12a2 and LwaCas13a, was treated as 100% activity, and the activity of each condition was standardized to this intensity.

## Results

### Apo, binary, and ternary Cas12a2 bind tightly to collateral ssRNA, ssDNA, and dsDNA substrates

Base pairing between the Cas12a2 crRNA-guide and a complementary target RNA drives a large conformational change in Cas12a2 that exposes and activates the RuvC nuclease active site (2) (**Figure 1A**). As substrate binding to the activated RuvC domain would be required for collateral cleavage, we reasoned that differences in affinity among collateral substrate types could reveal features of the collateral substrate cleavage mechanism, such as substrate preference. We also reasoned that apo (Cas12a2 alone) and binary complex (Cas12a2 bound to crRNA) would have some affinity for collateral nucleic acids, as apo binds the crRNA-guide within a positively charged cleft, and binary complex is primed to bind target RNA to form a ternary complex (Cas12a2, crRNA, and target RNA). To explore the binding events governing Cas12a2 activation, we determined the dissociation equilibrium constants (K_d_) of collateral substrates to apo, binary, and ternary complexes with fluorescence anisotropy, using FAM-labeled and non-hydrolysable phosphorothioated (PT) PT-ssRNA, PT-ssDNA, and PT-dsDNA substrates (**Figure 1C-E, Supplementary Figure S2 & Supplementary Table 1**). Binary and ternary complexes were made by adding RNAs (crRNA-guide or crRNA-guide and target RNA) in excess of Cas12a2, and then were SEC-purified to remove excess RNAs before adding them to binding assays.

**Figure 1.**
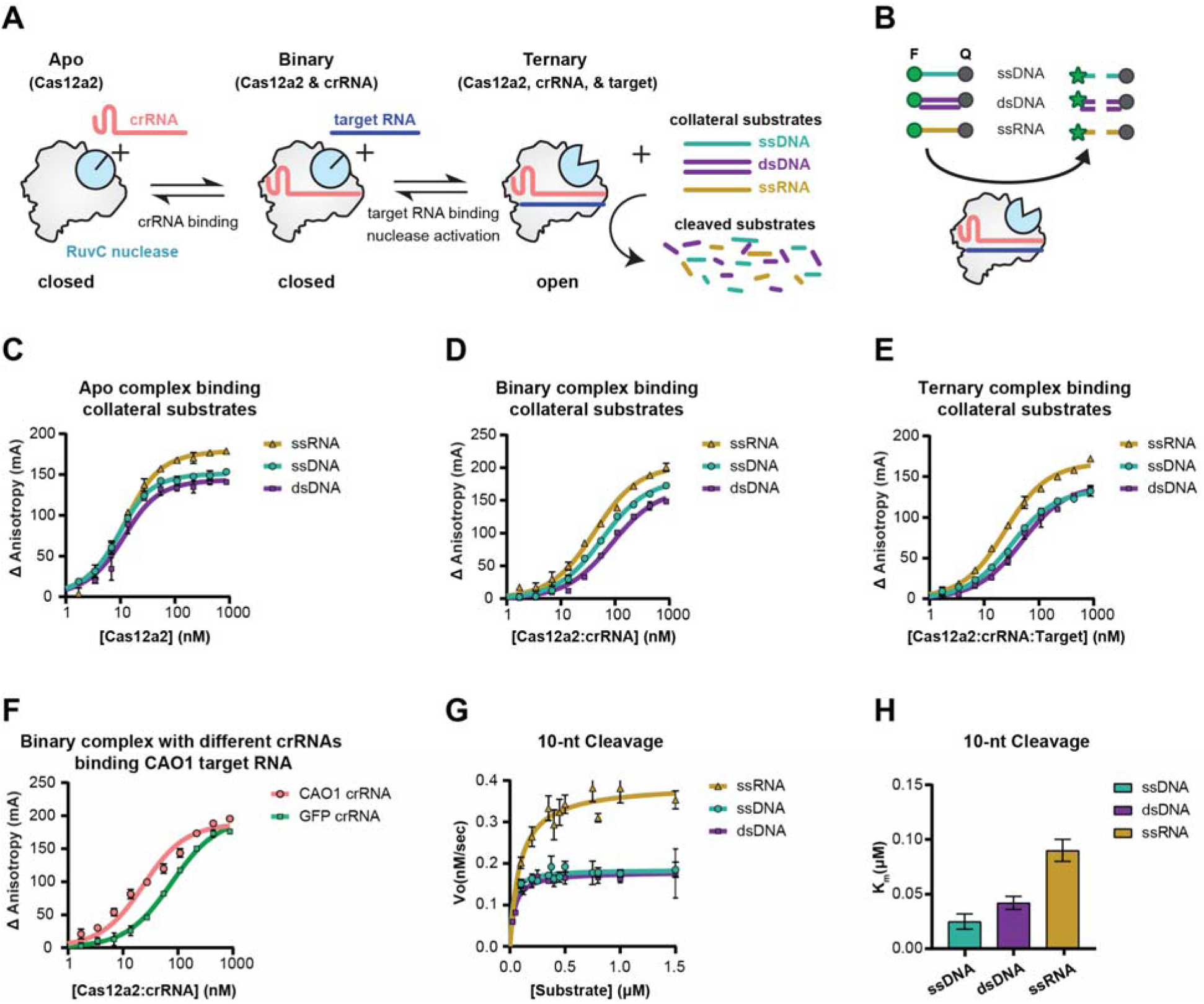
Cas12a2 preferentially cleaves collateral DNA substrates. **A.** Schematic representing the target RNA-triggered conformational changes that activate the Cas12a2 RuvC collateral nuclease activities leading to cleavage of ssRNA, ssDNA, and dsDNA. **B.** Schematic of reporter substrates with 5′ fluorescein (F) and 3′ Iowa Black quencher (Q) modifications used to measure Cas12a2 collateral cleavage activity. **C-E.** Fluorescence anisotropy binding isotherms of non-hydrolyzable, phosphorothioated (PT) collateral ssDNA (cyan), dsDNA (purple), and ssRNA (yellow) binding by Apo-Cas12a2 (C), Cas12a2 binary complex (D), and Cas12a2 ternary complex (E). **F.** Binding isotherms for non-hydrolyzable, phosphorothioated (PT) target RNA binding to binary complex with a complementary CAO1 crRNA (pink) and to binary complex with non-complementary GFP crRNA (green). **G.** Michaelis-Menten curves for 10-nt long collateral cleavage substrates. Data points represent data from at least two biological replicates collected in experimental triplicate. **H.** Comparison of K_m_ values for cleavage of different collateral substrates. Error bars show the standard deviation from the mean.

In line with the presence of a positively charged pocket that binds the crRNA, Apo Cas12a2 bound all three collateral substrates with at least a four-fold greater affinity than binary and ternary complexes with apparent Kds about 4-7 nM (**Figure 1C and Table 1**). Binary and ternary complexes also bound each collateral substrate with nanomolar affinities, with K_d_ values ranging from 37 to 82 nM for binary and 19 to 44 nM for ternary. As controls, we determined that bovine serum albumin (BSA) bound these same substrates in the 250-400 nanomolar range, and when crRNA and target RNA were combined without Cas12a2, no binding was detected, suggesting the measured binding was dependent on the presence of Cas12a2 (**Table 1and Supplemental Figure S3**). The ternary complex bound collateral substrates with about twice the affinity of the binary complex. Also, the ternary complex bound collateral ssRNA (Kd = 19 +/-1 nM) with the same affinity that the binary complex bound target RNA (Kd = 18 +/- 4 nM) (**Table 1**). These high affinities for collateral substrates of the ternary complex could be due to an increase in positive surface charge upon RNA-target-mediated opening of the RuvC nuclease domain (2). Both binary and ternary complexes bound collateral RNA with the highest affinity (binary Kd = 37 +/- 2 nM and ternary Kd = 19 +/- 1 nM), followed by ssDNA (binary Kd = 52 +/- 2 nM and ternary Kd =27 +/- 1 nM) and dsDNA (binary Kd = 82 +/- 5 nM and ternary Kd = 44 +/- 3 nM).

**Table 1.** Binding parameters determined by fluorescence anisotropy.

| **Table 1. Binding parameters determined by fluorescence anisotropy** |  |  |
| --- | --- | --- |
| **Substrate** | **Bmax (mA)** | **Apparent $K_d$ (nM)** |
| **Apo- Cas12a2** |  |  |
| Collateral ssRNA | 179 ± 2 | 6.7 ± 0.5 |
| Collateral ssDNA | 152 ± 1 | 4.1 ± 0.3 |
| Collateral dsDNA | 144 ± 3 | 6.1 ± 0.9 |
| **Cas12a2 binary complex (CAO1 crRNA)** |  |  |
| Target RNA | 189 ± 2 | 18 ± 4 |
| Collateral ssRNA | 203 ± 3 | 37 ± 2 |
| Collateral ssDNA | 182 ± 2 | 52 ± 2 |
| Collateral dsDNA | 168 ± 3 | 82 ± 5 |
| **Cas12a2 binary complex(GFP crRNA)** |  |  |
| Target RNA | 197 ± 2 | 72 ± 3 |
| **Cas12a2 ternary complex** |  |  |
| Collateral ssRNA | 167 ± 1 | 19 ± 1 |
| Collateral ssDNA | 134 ± 1 | 27 ± 1 |
| Collateral dsDNA | 140 ± 2 | 44 ± 3 |
| Nucleic acid substrates used are non-hydrolyzable, phosphorothioated (PT) substrates. |  |  |

We hypothesize that the higher affinity of binary Cas12a2 for collateral RNA over DNA substrates may be due to the binary conformation being primed for RNA target binding rather than DNA. However, complementarity to the crRNA-guide substantially increased affinity to binary Cas12a2, as a complementary target RNA bound with about two-fold greater affinity (target RNA to binary Kd = 18 +/- 4 nM (complementary CAO1 crRNA) compared to the non-complementary collateral ssRNA (Kd = 37 +/- 2 nM), and changing the crRNA guide sequence decreased affinity for the original target RNA (CAO1 target) about fourfold (Kd = 72 +/- 3 nM) **(Figure 1F and Table 1)**.

Affinity of non-complementary ssRNA for Cas12a2 binary complex appears to be sequence-dependent as non-complementary ssRNA added to Cas12a2-CAO1-gRNA (Kd = 37 +/- 2 nM), bound with about twice the affinity as target CAO1 RNA added to non-complementary Cas12a2-GFP-gRNA (Kd = 72 +/- 3 nM), even though both sequences have the same length (53 nt). The higher affinity of non-complementary ssRNA to the Cas12a2-CAO1-guide over the CAO1 target RNA to the Cas12a2-GFP-gRNA is consistent with differences in maximum delta G calculations for the guides and targets performed by IDT OligoAnalyzer tool (−6.21 and −4.71 kcal/mol, respectively). Therefore, we attribute the difference in affinity to the distinct guide and target sequences and their ability to form transient base pairs. Notably, the RNA-target affinities measured for Cas12a2 are comparable to those measured for other single-subunit RNA-targeting Cas nucleases, such as Cas13 (3, 19). Collectively, binding assays revealed that apo, binary, and ternary Cas12a2 complexes bind all collateral substrates with affinities in the sub-100-nanomolar range, with ssRNA binding ternary Cas12a2 with a higher affinity than ssDNA or dsDNA.

### Cas12a2 cleaves DNA substrates with greater catalytic efficiency

To more clearly understand the cleavage preferences of Cas12a2, we determined steady-state kinetic parameters for the cleavage of each collateral substrate **(Figure 1B)**. We hypothesized that we could measure initial velocities of collateral dsDNA, ssDNA, and ssRNA cleavage at increasing substrate concentrations using a fluorescent reporter assay, similar to previous work evaluating collateral cleavage activities of other single-subunit Cas nucleases (11). By fitting velocities to the Michaelis-Menten equation we determined the Michaelis constant (K_m_), the maximum reaction velocity (V_max_), and the catalytic rate constant (*k*_cat_) (See Methods), and with these parameters we compared activities using the catalytic efficiencies (*k*_cat_/K_m_) of cleavage, also known as specificity constants, for each collateral substrate (20, 21).

We reasoned that a 10-nt substrate would be long enough to bind correctly into the RuvC active site, given that 11 base pairs of a bound collateral dsDNA were resolved in the quaternary structure of Cas12a2 (2). Additionally, we reasoned that 10-nt would be short enough to limit Cas12a2 from cutting the probe more than once, allowing us to directly correlate increases in fluorescence with the concentration of cut probe. We chose a random, 50% GC content, nucleic acid sequence (5’-TGTAACGACC-3’ / 5’-UGUAACGACC-3’), without complementarity to the crRNA guide or target RNA sequences, to represent most collateral nucleic acid sequences. Using 10-nt probes of each collateral substrate type, each with the same sequence (**Supplemental Table 1**), and an SEC-purified ternary Cas12a2 complex, we observed robust cleavage toward ssRNA, ssDNA, and dsDNA at various concentrations (**Figure 1G**, **Table 2, and Supplemental Figure S4-1**). Fitting these velocities to the Michaelis-Menten equation indicated the collateral ssRNA probe is cleaved with a 2-fold larger turnover number (*k*_cat_ = 0.0260 +/- 0.0007 sec^-1^) than for the ssDNA (*k*_cat_ = 0.0123 +/- 0.0003 sec^-1^) and dsDNA (*k*_cat_ = 0.0120 +/- 0.0003 sec^-1^) probes. The higher ssRNA probe cleavage turnover number is consistent with the larger V_max_ (0.39 +/- 0.01 nM sec^-1^) and K_m_ (0.09 +/- 0.01 μM) values for ssRNA probe cleavage than for ssDNA (V_max_ = 0.185 +/- 0.004 nM sec^-1^, K_m_ = 0.025 +/- 0.007 μM) and dsDNA probe cleavage (V_max_ = 0.180 +/- 0.005 nM sec^-1^, K_m_ = 0.042 +/- 0.006 μM) (**Figure 1G and 1H and Table 2**).

**Table 2.** Cas12a2 collateral cleavage kinetics.

| Table 2. Cas12a2 collateral cleavage kinetics. |  |  |  |  |  |  |
| --- | --- | --- | --- | --- | --- | --- |
| Substrate | $K_m$ ($\mu M$) | | $V_{max}$ ($nM\ sec^{-1}$) | | $k_{cat}$ ($sec^{-1}$) | $k_{cat}/K_m$ ($M^{-1}\ sec^{-1}$) |
| 10-nt Collateral Substrates |  |  |  |  |  |  |
| ssRNA | 0.09 | $\pm 0.01$ | 0.39 | $\pm 0.01$ | $0.0260 \pm 0.0007$ | $(2.9 \pm 0.3) \times 10^5$ |
| 5'-UGUAACGACC-3' |  |  |  |  |  |  |
| Cas12a2 (Y465A) | 0.104 | $\pm 0.008$ | 0.422 | $\pm 0.009$ | $0.0281 \pm 0.0006$ | $(2.7 \pm 0.2) \times 10^5$ |
| Cas12a2 (Y1080A) | 0.12 | $\pm 0.01$ | 0.359 | $\pm 0.009$ | $0.0239 \pm 0.0006$ | $(2.0 \pm 0.2) \times 10^5$ |
| Cas12a2 (Y1069A) | 0.16 | $\pm 0.01$ | 0.58 | $\pm 0.01$ | $0.0387 \pm 0.0007$ | $(2.4 \pm 0.2) \times 10^5$ |
| Cas12a2 (F1092A) | 3.6 | $\pm 3.7$ | 0.5 | $\pm 0.4$ | $0.03 \pm 0.03$ | $(1.0 \pm 1.7) \times 10^4$ |
| ssDNA | 0.025 | $\pm 0.007$ | 0.185 | $\pm 0.004$ | $0.0123 \pm 0.0003$ | $(4.9 \pm 1.4) \times 10^5$ |
| 5'-TGTAACGACC-3' |  |  |  |  |  |  |
| Cas12a2 (Y465A) | 0.025 | $\pm 0.004$ | 0.243 | $\pm 0.005$ | $0.0162 \pm 0.0003$ | $(6.5 \pm 1.1) \times 10^5$ |
| Cas12a2 (Y1080A) | 0.015 | $\pm 0.003$ | 0.189 | $\pm 0.004$ | $0.0126 \pm 0.0003$ | $(8.4 \pm 1.7) \times 10^5$ |
| Cas12a2 (Y1069A) | 0.030 | $\pm 0.007$ | 0.251 | $\pm 0.009$ | $0.0167 \pm 0.0006$ | $(5.6 \pm 1.3) \times 10^5$ |
| Cas12a2 (F1092A) | 0.25 | $\pm 0.06$ | 0.054 | $\pm 0.004$ | $0.0036 \pm 0.0003$ | $(1.4 \pm 0.4) \times 10^4$ |
| dsDNA | 0.042 | $\pm 0.006$ | 0.18 | $\pm 0.005$ | $0.0120 \pm 0.0003$ | $(2.9 \pm 0.4) \times 10^5$ |
| 5'-F-TGTAACGACC-Q-3' |  |  |  |  |  |  |
| 3'- ACATTGCTGG -5' |  |  |  |  |  |  |
| Cas12a2 (Y465A) | 0.10 | $\pm 0.03$ | 0.18 | $\pm 0.01$ | $0.0120 \pm 0.0007$ | $(1.2 \pm 0.4) \times 10^5$ |
| Cas12a2 (Y1080A) | 0.03 | $\pm 0.01$ | 0.14 | $\pm 0.01$ | $0.0093 \pm 0.0007$ | $(3.1 \pm 1.1) \times 10^5$ |
| Cas12a2 (Y1069A) | 0.10 | $\pm 0.02$ | 0.19 | $\pm 0.01$ | $0.0127 \pm 0.0007$ | $(1.3 \pm 0.3) \times 10^5$ |
| Cas12a2 (F1092A) | -- |  | -- |  | -- | -- |
| dsDNA( Quencher on opposite strand) | 0.040 | $\pm 0.005$ | 0.218 | $\pm 0.005$ | $0.0145 \pm 0.0003$ | $(3.6 \pm 0.5) \times 10^5$ |
| 5'-F-TGTAACGACC -3' |  |  |  |  |  |  |
| 3'- ACATTGCTGG-Q-5' |  |  |  |  |  |  |

The turnover numbers for ssDNA and dsDNA probe cleavage are approximately the same (**Table 2**). Assuming the non-labeled strand of the duplex has the same probability as the labeled strand to dock into the RuvC active site, this result revealed the possibility that at saturating conditions, Cas12a2 may cleave the two strands of the dsDNA probe in the same time period that it cleaves the ssDNA probe. To rule out the possibility that the similar turnover numbers are due to preferential binding of the FAM-labeled strand of the dsDNA probe into the RuvC active site, we designed a new dsDNA probe of the same sequence, but with the FAM and quencher each on the 5′ ends of opposite cognate strands of the dsDNA probe. Under this design, cutting one strand would release the FAM label, while cutting the other strand would release the quencher, enabling detection of single cuts within the dsDNA probe. Cas12a2 cleaved this second dsDNA probe with a slightly higher turnover number (**Table 2 and Supplemental Figure S6**), consistent with a mechanism where Cas12a2 makes single-strand cuts (nicks) in dsDNA probes more often than it cleaves both strands. Notably, our plasmid cleavage data, described below, also support such a mechanism.

Although Cas12a2 cleaves the ssRNA probe with a turnover number twice that for ssDNA and dsDNA probes, the larger K_m_ for ssRNA probe cleavage compared to ssDNA and dsDNA probe reduced the specificity constant (*k*_cat_/K_m_ = (2.9 +/- 0.3) × 10L M^-1^ sec^-1^) to about 60% that of ssDNA cleavage (*k*_cat_/K_m_ = (4.9 +/- 1.4) × 10L M^-1^ sec^-1^) and to a similar value of the specificity constant for cleavage of the dsDNA probe with the FAM and quencher on the same strand (*k*_cat_/K_m_ = (2.9 +/- 0.4) × 10L M^-1^ sec^-1^). Additionally, the specificity constant for cleaving the other dsDNA probe labeled with the FAM and quencher on opposite strands (*k*_cat_/K_m_ = (3.6 +/-0.5) × 10L M^-1^ sec^-1^) is slightly greater than that for the ssRNA probe, suggesting that Cas12a2 nicks the dsDNA probe with greater efficiency than cleaving the ssRNA probe. Quantitatively, these kinetic parameters differ from those determined previously with Cas12a2 and collateral substrates of different sequences and lengths (16), indicating that sequence and length of collateral substrates influence kinetic parameters. However, qualitatively, the values determined here are within a similar range of previously determined Cas12a2 specificity constants (10^4^ - 10^5^ M^-1^ sec^-1^) and ssDNA was preferably cleaved over ssRNA in both studies (**Supplementary Table 4**) (16). Additionally, Cas12a2 cleaves collateral ssDNA with similar activity to other type V nucleases (Cas12a1 = 10^4^ - 10^6^ M^-1^ sec^-1^), but with about 10-fold smaller ssRNA collateral cleavage than type VI nucleases (Cas13 = 10^6^ M^-1^ sec^-1^) (**Supplementary Table 4**) (11).

The specificity constant for cleavage of the dsDNA probe with a FAM and a quencher on the same strand is about 60% smaller than for ssDNA probe cleavage, while the other dsDNA probe is 73% smaller, but within error (**Table 2**). The smaller specificity constants for the cleavage of dsDNA probes are due to the two-fold larger K_m_ values for dsDNA cleavage compared to ssDNA cleavage. As this difference is consistent with the two-fold lower affinity measured for dsDNA compared to ssDNA in the ternary complex, the lower specificity constant may be attributed to thermodynamic penalties associated with bending and unwinding duplex DNA to properly bind to the RuvC active site in a conformation amenable to cleavage (2). On the other hand, the ternary complex binds ssRNA with greater affinity than ssDNA and dsDNA. Thus, the larger K_m_ for ssRNA probe cleavage cannot be explained solely by differences in affinity. Collectively, these data indicate that Cas12a2 prefers to cleave DNA over RNA, as it cleaves ssDNA with the greatest catalytic efficiency (i.e., the greatest specificity constant) and cleaves dsDNA with superior or equivalent efficiency to RNA, depending on probe labeling.

### Cas12a2 preferentially cleaves AT-rich sequences in plasmid DNA

We and others previously showed that activated Cas12a2 rapidly degrades supercoiled plasmid DNA (1, 2, 14). However, it remained unclear how negative supercoiling would affect dsDNA cleavage and whether target RNA-triggered *Sulfuricurvum* sp. PC08-66 (Su)Cas12a2 prefers to cleave one dsDNA sequence over another. To better understand the cut-site preference of Cas12a2 for supercoiled dsDNA, we activated RNA-guided Cas12a2 with a complementary target and incubated it with supercoiled pUC19. To identify the Cas12a2-mediated cut sites, we gel-purified pUC19 plasmids linearized by Cas12a2 cleavage, ligated next-generation sequencing primers to the 5′ ends of the linear fragments, and performed next-generation sequencing (**Figure 2A**). As a control, we also sequenced pUC19 linearized with the restriction enzyme BamHI, which targets a single site within pUC19 (**Figure 2B**). As expected, sequencing results revealed a single distinct cut site for the BamHI control, while Cas12a2 cleavage sites decorated the entire plasmid, with one site within the ampicillin promoter appearing two to four times more often than most other cut sites (**Supplemental Table 2**). This sequence contains a span of AT base pairs, consistent with a preference to cleave dsDNA that is more easily melted. To explore whether a similar sequence preference occurred at the other Cas12a2 cut sites, we identified sequences within a 14-bp window around each cut site and analyzed them using WebLogo (18) (**Figure 2C**, **Supplemental Table 2**). While no conserved cleavage motif emerged, even when excluding the preferred cut-site (**Figure 2C bottom**), we noticed a slightly higher probability of adenine or thymine around the cut site than guanine and cytosine.

**Figure 2.**
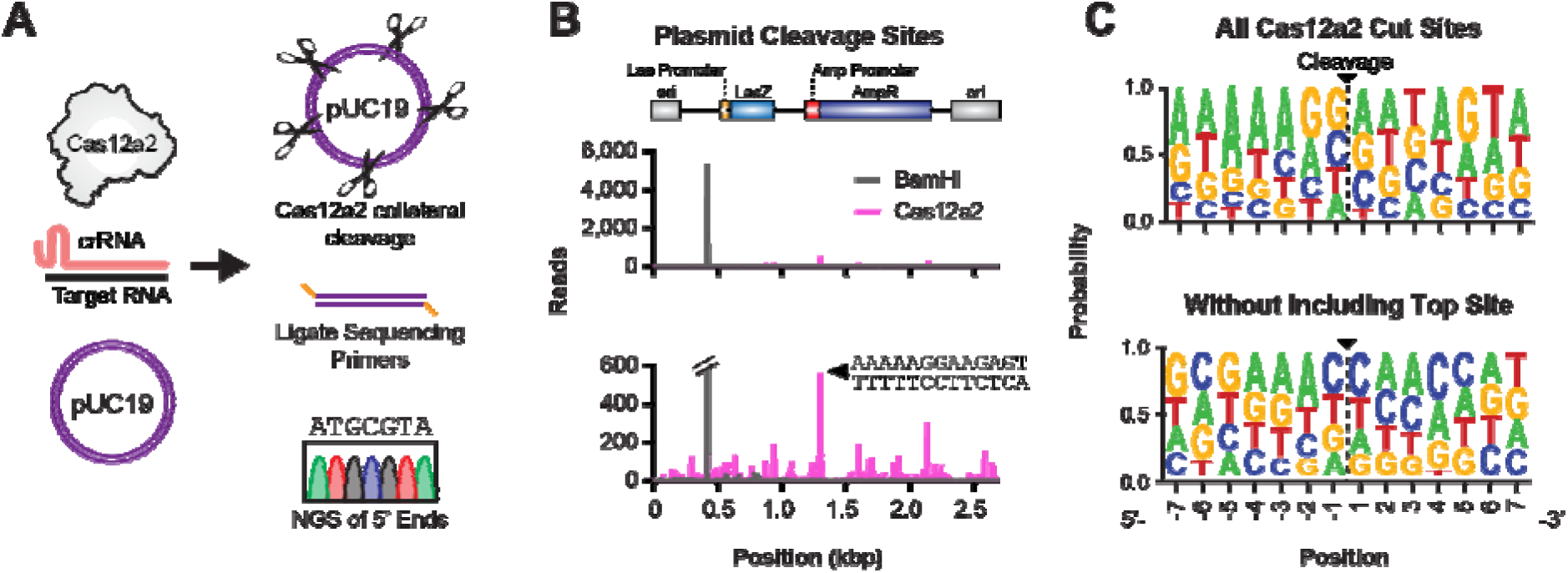
Cas12a2 preferentially cleaves AT-rich sequences in plasmid DNA. **A.** Schematic of the method used to determine Cas12a2 cut-sites in supercoiled plasmid DNA. **B.** Histogram of pUC19 plasmid cut-sites identified by next-generation sequencing, with number of reads on the vertical-axis and plasmid position on the horizontal-axis. The annotated pUC19 sequence is shown at the top. **C.** WEBLOGO of flanking sequences around all Cas12a2 cut-sites (top), and Cas12a2 cut-sites with the top site sequence not included in the analysis. Cas12a2 cuts between position 1 and −1 as indicated.

However, the presence of extended spans of AT base pairing was not a requirement for plasmid cleavage (**Supplemental Table 2**). To fit dsDNA into the active site, Cas12a2 bends dsDNA 90° and melts two base pairs, including the base containing the scissile phosphate (2). Therefore, we hypothesize that AT-rich sequences are more represented in our plasmid cleavage experiment because they are more easily distorted than GC-rich sequences. Together, these plasmid cleavage data suggest that dsDNA sequences with lower melting temperatures correlate with higher Cas12a2 cleavage efficiencies.

### Aromatic clamps contribute to DNA cleavage

The melting and bending of collateral dsDNA into the Cas12a2 RuvC nuclease active site was previously proposed to depend on a system of aromatic clamps (Y465, Y1069, Y1080, F1092) that interact directly with collateral dsDNA in the RuvC nuclease active site and are essential for *in vivo* function (**Figure 3A-B and Supplemental Figure S7**) (2). Residues Y1069 and F1092 pi-stack, or ‘clamp’, the sides of a nucleobase located within the ‘cleaved strand’ of the duplex DNA, positioning the adjacent scissile phosphate into the RuvC active site, while residues Y465 and Y1080 pi-stack the complementary nucleotide of the ‘non-cleaved strand’ (**Figure 3B**).

**Figure 3.**
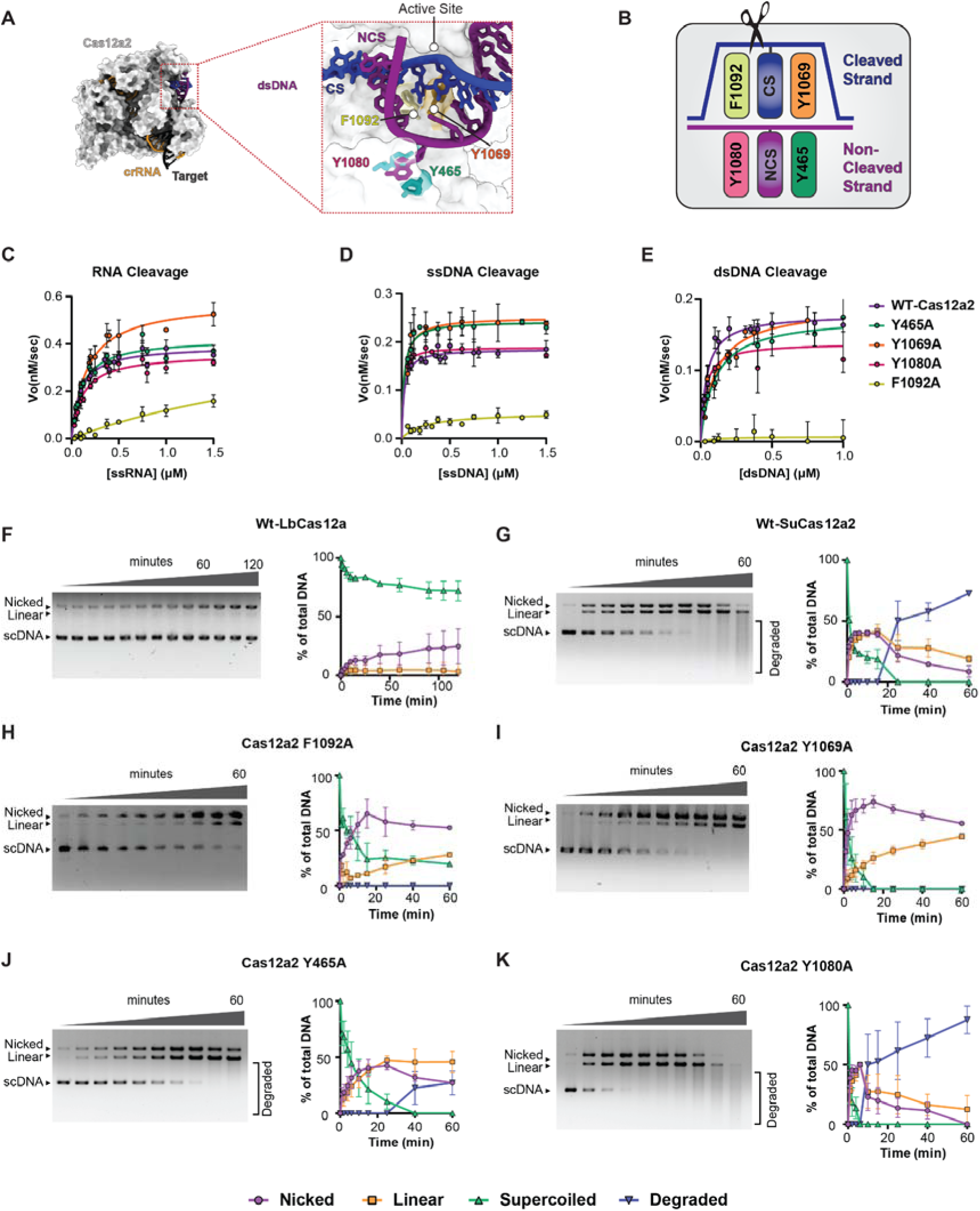
The aromatic clamps contribute to DNA cleavage. **A.** Cryo-EM structure (PDBid 8D4A) of Cas12a2 bound to crRNA, target RNA, and collateral dsDNA (quaternary structure). The cleavage stran (CS) and non-cleavage strand (NCS) are each coordinated by a pair of aromatic clamps (inset). **B.** Schematic of which aromatic clamps coordinate which strand of DNA. Scissors indicate the nuclease active site. **C-E.** Michaelis-Menten kinetics of cleavage of ssRNA (C), ssDNA (D), and dsDNA (E) by wild-type (WT) Cas12a2 compared to the aromatic clamp mutants. Wild-type cleavage curves represent replotting of data previously shown in Figure 1G **F-K.** Plasmid (pUC19) cleavage assays. Quantification plots with corresponding agarose gels are shown for wild-type LbCas12a1 (F), wild-type Cas12a2 (G), and aromatic clamp mutants F1092A (H), Y1069A (I), Y465A (J), and Y1080A (K).

Point mutants of the aromatic residues previously showed that each was important for dsDNA cleavage and Cas12a2-mediated immunity in bacteria (2). Since Y465A, Y1080A, and Y1069A were still robustly able to cleave ssDNA and ssRNA, it was concluded that the immunity defects caused by mutating the aromatic clamps were due to impaired dsDNA cleavage (2). Since the aromatic clamps appeared to be involved in binding and positioning dsDNA substrates for cleavage, we hypothesized that they would be involved in the mechanism of cleaving both strands of collateral dsDNA substrates.

To test the impact of the aromatic clamp mutations on collateral substrate cleavage, we expressed and purified ternary complexes of Cas12a2 mutants F1092A, Y1069A, Y465A, and Y1080A and determined the steady-state kinetic parameters of collateral ssRNA, ssDNA, and dsDNA cleavage (**Figure 3C-E and Supplemental Figure S8**). As previously observed (2), the F1092A mutation impaired cleavage of all three substrates, while Y1069A and Y465A only disrupted dsDNA cleavage. In contrast to our previous work, Y1080A demonstrated no negative effect on the cleavage of any collateral substrate, including dsDNA (**Table 2**). The dsDNA cleavage activity of F1092A was so reduced that measured velocities would not fit the Michaelis-Menten equation, precluding calculation of kinetic parameters. For ssDNA and ssRNA cleavage, the K_m_ values for F1092A were ten- and four-fold greater than wild type, respectively, and the turnover numbers for ssDNA and ssRNA cleavage were respectively four-fold smaller, or similar to wild type, but with a large error. The large K_m_ values for ssDNA and ssRNA, combined with the smaller or equivalent turnover numbers, resulted in specificity constants for ssDNA and ssRNA cleavage approximately thirty-fold smaller than wild type for the F1092A mutant. These defects are consistent with the F1092A mutant causing the largest immune system defect *in vivo*, and the Cas12a2 quaternary structure, which shows F1092 helping position the scissile phosphate of dsDNA substrates into the RuvC nuclease active site (2).

Although previous work showed that Y1069A, Y465A, and Y1080A mutations impair Cas12a2-mediated immunity, all three mutants cleaved collateral ssRNA and ssDNA with kinetics similar to, or slightly faster than, wild type (**Table 2**, **Supplemental Figure S8**). Thus, unlike F1092, these aromatic residues are not critical for positioning or cleaving single-stranded nucleic acid substrates. On the other hand, the Y1069A (cleaved strand binding site) and Y465A (non-cleaved strand binding site) mutants impaired cleavage of collateral dsDNA, each causing a two-fold increase in Km, correlating with a two-fold reduction in the specificity constant. (**Table 2**, **Supplemental Figure S8**). In contrast, the Y1080A mutant showed no kinetic defects with any of the collateral substrates we tested, including dsDNA. While this mutation previously had the smallest impact on Cas12a2 immunity of all the aromatic clamp mutants (2), we were surprised to observe no kinetic defects beyond a slightly lower, but within error, turnover number compared to wild type. Collectively, these results strongly support a role for F1092 in positioning all collateral nucleic acid substrates into the RuvC active site, with Y465 and Y1069 involved in stabilizing dsDNA in a conformation that allows cleavage, and Y1080 playing a less important role in the cleavage mechanism across all collateral substrate types.

### Cas12a2 relies on aromatic clamp residues to degrade supercoiled DNA

Our kinetic results with the aromatic clamp mutants suggested that SuCas12a2 interacts with and cleaves dsDNA differently from single-stranded substrates. Considering these data and the similar wild-type turnover numbers for cleavage of collateral ssDNA and dsDNA, we hypothesized that the aromatic clamps may play a role in a mechanism that cleaves both strands of the duplex substrate in the same amount of time as it takes to cleave a single-stranded substrate, when the enzyme is under fully saturated conditions. However, we were concerned that our small dsDNA probe might not fully represent a natural double-stranded cleavage substrate for Cas12a2, as fraying of the ends of the DNA duplex could expose ssDNA to the active site for cleavage instead of fully duplexed dsDNA. Indeed, previous studies developing Cas12a1 as a diagnostic tool that observed dsDNA-triggered Cas12a1 could cleave small DNA probes with duplex features (6, 7, 11, 12, 10, 8, 9, 22), hypothesized dsDNA probe cleaving activity was due to fraying of the duplex ends or to available end overhangs (23, 12, 13, 10). Thus, to examine the collateral dsDNA mechanism of SuCas12a2 with a more natural substrate, we decided to measure the cleavage of closed-circle supercoiled plasmid over time, which would report single-stranded breaks as nicked or ‘open circle’ plasmid, double-stranded breaks as linear, and multiple double-stranded breaks as a degraded smeared band.

To determine whether Cas12a1 and Cas12a2 cleave dsDNA differently, we first compared plasmid cleavage activities of equivalent concentrations of target RNA-triggered SuCas12a2 and target DNA-triggered LbCas12a1. We used a supercoiled pUC19 plasmid with no complementarity to the Cas12a1 or Cas12a2 guide. Consistent with what has been previously reported (13), ternary LbCas12a1 quickly nicked some of the supercoiled plasmid, but even after 120 minutes, most of the plasmid remained uncleaved in the supercoiled fraction, and only a small percentage was found in the linear fraction (**Figure 3F**). The ability of Cas12a1 to nick some but not all of the dsDNA plasmid is consistent with supercoiled pUC19 locally unwinding to form ssDNA ‘bubbles’ that the indiscriminate ssDNAse activity of Cas12a1 could cleave (24). However, closed-circle dsDNA appears not to be the collateral substrate of Cas12a1, as most of the supercoiled plasmid remained uncleaved over time, and the small linear fraction that appeared at later time points could be attributed to the buildup of multiple single-stranded nicking events in the nicked fraction. In contrast to the LbCas12a1 product profile, SuCas12a2 activity produced equal fractions of linear and nicked DNA from the first 1-minute time point until the nicked fraction was depleted and replaced by linear or degraded bands. The equal portions of linear and nicked products indicate that about 50% of the time SuCas12a2 makes double-stranded breaks in collateral supercoiled DNA, while nicking DNA the other 50% of the time (**Figure 3G**).

Additionally, in contrast to the inability of LbCas12a1 to degrade all of the supercoiled plasmid within 120 minutes, target RNA-triggered SuCas12a2 cleaved all supercoiled plasmid into nicked or linear fractions within 25 minutes, and greater than 50% of the plasmid was degraded into a smeared band by 60 minutes (**Figure 3G**). Similar activity was previously observed using a plasmid cleavage assay to confirm Cas12a2 activity (1, 16).

To further investigate the Cas12a2 collateral cleavage mechanism of dsDNA, we examined how the aromatic clamp mutants cleave supercoiled plasmid. Consistent with our kinetics data, F1092A demonstrated the greatest cleavage defect, followed by Y1069A and Y465A, with Y1080A cleaving the plasmid similarly to wild type. Analyzing the cleavage profiles, we observed that mutating residues that interact with the cleaved strand (F1092A and Y1069A) caused SuCas12a2 to behave more like a nickase than wild-type Cas12a2. Cleavage products from these mutants populated the nicked fraction well before the linear fraction; the nicked fraction stayed more abundant than the linear fraction until the supercoiled fraction was fully depleted; and at 60 minutes, there were no smeared bands to indicate double-stranded breaks of the linear fraction (**Figure 3H-I**). In contrast, the cleavage product profile of Y465A was more similar to wild-type, producing equal percentages of nicked and linear DNA, and eventually a smeared DNA band, although the overall cleavage activity was reduced compared to wild type (**Figure 3J-K**). Collectively, these data suggest the aromatic residues that engage with the cleaved strand (F1092 and Y1069) are important for cleaving both strands of duplex DNA, while the aromatic residues that engage with the non-cleaved strand (Y465 and Y1080) are likely more involved in stabilizing the melted and bent conformation of dsDNA.

### Pairwise competition assays confirm a cleavage preference for collateral DNA

In bacterial systems, there is ∼7-fold more RNA than DNA in the cell cytoplasm (25). Comparison of collateral substrate specificity constants suggested that ssDNA is the preferred collateral cleavage substrate of Cas12a2. However, the higher cleavage turnover number and higher affinity for ssRNA, and fairly equal specificity constants for ssRNA and dsDNA cleavage, left it unclear how the presence of competing collateral substrates might affect Cas12a2 collateral cleavage activities. To explore the competition between ssRNA, ssDNA, and dsDNA substrates, we designed pairwise competition assays between ssRNA, ssDNA, and dsDNA with non-cleavable phosphorothioated PT-ssDNA or PT-RNA, as inhibitors (**Figure 4A**).

**Figure 4.**
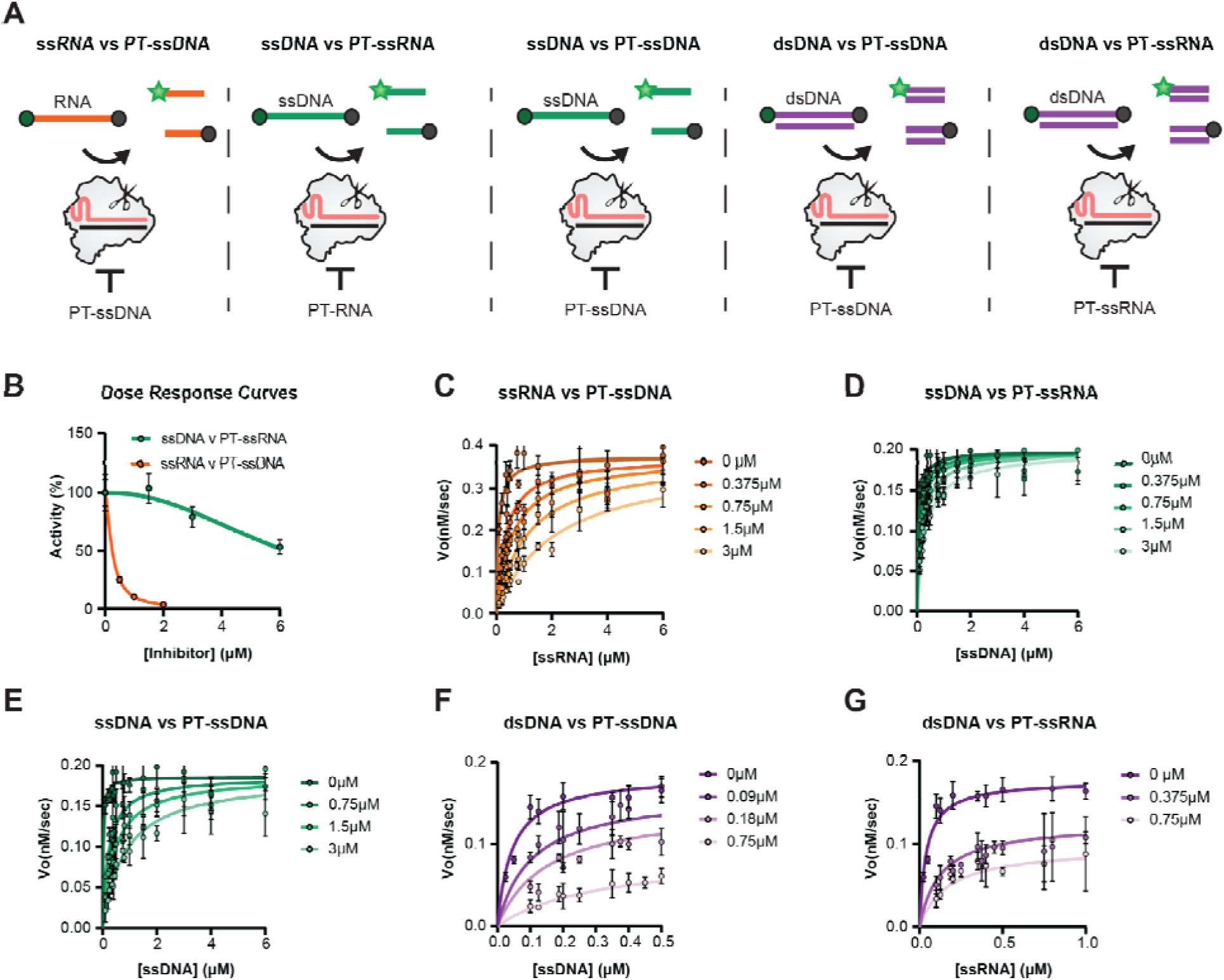
Inhibition kinetics of collateral substrate cleavage mechanisms. **A.** Schematic of pairwise competition assays of RNA cleavage with PT-ssDNA inhibition, ssDNA cleavage with PT-RNA inhibition, ssDNA cleavage with PT-ssDNA inhibition, dsDNA cleavage with PT-ssDNA inhibition, and dsDNA cleavage with PT-ssRNA inhibition, from left to right. **B.** Dose response curves for collateral ssDNA cleavage inhibition by PT-RNA (green) and collateral ssRNA cleavage inhibition by PT-ssDNA (orange). Data are represented as a percent of uninhibited initial velocity of 1µM RNA or ssDNA substrate cleavage. **C.** ssRNA cleavage kinetic plots showing the impact of increasing concentrations of PT-ssDNA on activity. **D.** ssDNA cleavage kinetic plots showing the impact of increasing PT-ssRNA concentration on activity. **E.** ssDNA cleavage kinetic plots showing the impact of increasing concentrations of PT-ssDNA on activity. **F.** dsDNA cleavage kinetic plots showing the impact of increasing concentrations of PT-ssDNA on activity. **G.** dsDNA cleavage kinetic plots showing the impact of increasing concentrations of PT-ssRNA on activity. The 0 µM curves of panels C-G represent the replotting of the cleavage data previously shown in Figure 1G.

We first examined ssRNA and ssDNA cleavage inhibition by PT-ssDNA and PT-ssRNA by generating inhibitor dose-response curves using a single concentration of cleavable collateral substrate and increasing concentrations of competing PT-substrate. We fit the data to the Michaelis-Menten equation to determine the type of inhibition and inhibition constants (K_i_) (**Figure 4B**). We found that increasing concentrations of PT-ssDNA drastically reduced ssRNA cleavage (Ki = 0.12 +/- 0.01 µM), whereas increasing PT-ssRNA concentrations weakly affected ssDNA cleavage (Ki = 0.61 +/- 0.09 μM). The inhibition of PT-ssDNA on ssRNA cleavage was ∼ six-fold stronger than the inhibition of ssDNA cleavage by PT-RNA (**Figure 4C-D and Table 3**), consistent with ssDNA as the preferred cleavage substrate.

**Table 3.** Kinetic parameters of pairwise competiton experiments.

| Table 3. Kinetic parameters of pairwise competition experiments |  |  |  |
| --- | --- | --- | --- |
| Substrate | $K_m$ ($\mu\text{M}$) | | $V_{\max}$ ($\text{nMsec}^{-1}$) |
| ssRNA vs PT-ssDNA ($K_i = 0.12 \pm 0.01\mu\text{M}$) [Competitive] | | | |
| 0 | 0.09 | $\pm 0.01$ | 0.39 $\pm 0.01$ |
| 0.375 | 0.33 | $\pm 0.03$ | 0.364 $\pm 0.009$ |
| 0.75 | 0.55 | $\pm 0.04$ | 0.36 $\pm 0.01$ |
| 1.5 | 1.5 | $\pm 0.2$ | 0.46 $\pm 0.02$ |
| 3.0 | 2.7 | $\pm 0.3$ | 0.44 $\pm 0.02$ |
| ssDNA vs PT-ssRNA ($K_i = 0.61 \pm 0.09\mu\text{M}$) [Competitive] | | | |
| 0.0 | 0.025 | $\pm 0.007$ | 0.185 $\pm 0.004$ |
| 0.375 | 0.09 | $\pm 0.01$ | 0.199 $\pm 0.004$ |
| 0.75 | 0.18 | $\pm 0.02$ | 0.209 $\pm 0.005$ |
| 1.5 | 0.21 | $\pm 0.02$ | 0.196 $\pm 0.004$ |
| 3.0 | 0.34 | $\pm 0.02$ | 0.214 $\pm 0.005$ |
| ssDNA vs PT-ssDNA($K_i = 0.09 \pm 0.02\mu\text{M}$)[Competitive] | | | |
| 0.0 | 0.025 | $\pm 0.007$ | 0.185 $\pm 0.004$ |
| 0.75 | 0.24 | $\pm 0.04$ | 0.196 $\pm 0.008$ |
| 1.5 | 0.46 | $\pm 0.06$ | 0.188 $\pm 0.007$ |
| 3.0 | 0.71 | $\pm 0.07$ | 0.169 $\pm 0.006$ |
| dsDNA vs PT-ssDNA ($K_i = 0.05 \pm 0.01\mu\text{M}$ and $K_i' = 0.7 \pm 0.2\mu\text{M}$) [Mixed] | | | |
| 0.0 | 0.042 | $\pm 0.006$ | 0.180 $\pm 0.005$ |
| 0.09375 | 0.09 | $\pm 0.02$ | 0.17 $\pm 0.01$ |
| 0.1875 | 0.20 | $\pm 0.03$ | 0.147 $\pm 0.007$ |
| 0.75 | 0.27 | $\pm 0.07$ | 0.087 $\pm 0.009$ |
| dsDNA vs PT-ssRNA ($K_i = 0.11 \pm 0.03\mu\text{M}$ and $K_i' = 0.9 \pm 0.2\mu\text{M}$) [Mixed] | | | |
| 0.0 | 0.042 | $\pm 0.006$ | 0.180 $\pm 0.005$ |
| 0.375 | 0.11 | $\pm 0.03$ | 0.114 $\pm 0.007$ |
| 0.75 | 0.17 | $\pm 0.04$ | 0.107 $\pm 0.009$ |

Both PT-ssRNA and PT-ssDNA appear to act as respective competitive inhibitors toward ssDNA and ssRNA cleavage, as increasing the inhibitor concentration effectively raised the K_m_ value, while the V_max_ remained approximately the same **(Supplemental Figure S9)**. Also, in support of a competitive inhibition mechanism, increasing concentrations of PT-ssDNA to the ssDNA cleavage reaction also increased the K_m_ without changing Vmax, and yielded a similar Ki (0.09 +/- 0.02 µM) to that calculated for inhibition of ssRNA cleavage (Ki = 0.12 +/- 0.01 µM) (**Figure 4E**).

PT-ssDNA and PT-ssRNA also inhibited the cleavage of our dsDNA probe. When fit to the Michaelis-Menten equation, increasing concentrations of either inhibitor increased the K_m_ and decreased the V_max_, suggesting that PT-ssDNA and PT-ssRNA act as mixed inhibitors. The mixed inhibition model indicates that (i) the inhibitors compete with the dsDNA probe for binding to ternary Cas12a2 RuvC active site and (ii) impair cleavage by binding to the ternary Cas12a2-dsDNA probe complex. Two different constants describe the inhibition, with K_i_ representing the ratio of [E][I] / [EI], E = activated Cas12a2, [EI] = activated Cas12a2 bound to the inhibitors, and I = the inhibitors, and K_i_′ representing the ratio [ES][I]/[ESI] where S = the fluorescently labeled dsDNA probe (**Figure 4F-G and Table 3**). Fitting the inhibition curves to the mixed inhibition equation revealed that PT-ssDNA inhibited dsDNA cleavage with a 2-fold smaller K_i_ (0.05 +/-0.01 µM) than for ssDNA (K_i_ = 0.09 +/- 0.02) or ssRNA (K_i_ = 0.12 +/- 0.01 µM) cleavage.

Likewise, PT-ssRNA inhibited dsDNA cleavage with a much smaller K_i_ (0.11 +/- 0.03 µM) (greater inhibition) than for ssDNA (K_i_ = 0.61 +/- 0.09 µM). The greater inhibition of dsDNA cleavage is consistent with the single-stranded inhibitor binding either the cleaved strand or the non-cleaved strand binding site, thereby impairing dsDNA cleavage activity. Additionally, PT-ssDNA inhibits dsDNA cleavage with a K_i_ half that of PT-ssRNA, indicating that PT-ssDNA is the stronger inhibitor, consistent with inhibition trends for single-stranded probe cleavage. In contrast to the differing K_i_ values, the K_i_′ values of PT-ssRNA and PT-ssDNA toward dsDNA cleavage were more similar (PT-ssRNA K_i_′ = 0.9 +/- 0.2 µM, and PT-ssDNA K_i_′ = 0.7 +/- 0.2 µM).

### Cas12a2 specificity for DNA substrates enhances RNA detection

RNA extraction from cells for the purposes of RNA diagnostics necessarily involves the purification of many non-target RNAs. We hypothesized that the collateral cleavage of fluorescent RNA probes by RNA-targeting CRISPR nucleases such as Cas13a might be hindered by competing binding and cleavage between the fluorescent RNA probe and non-target RNAs. Additionally, in scenarios where pre-amplification of on-target RNAs is performed, cleavage competition between the collateral RNA probe and excess target RNA could reduce the signal strength. After discovering that RNA-activated Cas12a2 has a strong preference for collateral ssDNA over ssRNA (i.e., PT-ssRNA did not strongly inhibit ssDNAse cleavage), we reasoned that Cas12a2, in combination with a ssDNA probe, would not face the same level of competition between probe and non-target substrates.

To test the impact of non-target RNA concentration on probe cleavage, we desired to compare the endpoint fluorescence signals of Cas12a2 cleaving a ssDNA probe and *Leptotrichia wadei* (Lwa)Cas13a cleaving an RNA probe in the presence of competing total yeast RNA (**Figure 5A**). The optimal endpoint reporter sequence for Cas13a was previously found to be polyuridine substrates (26). To identify the best endpoint reporter sequence for Cas12a2, we systematically screened 6-nt-long dinucleotide DNA triplet sequence combinations (**Supplemental Figure S10**). We observed the highest fluorescence signal over background with the T6 ssDNA reporter substrate, indicating that it is an optimal sequence for an RNA detection probe and consistent with the T-rich 5-mer probe used by another group to develop Cas12a2 as an RNA diagnostic (16).

**Figure 5.**
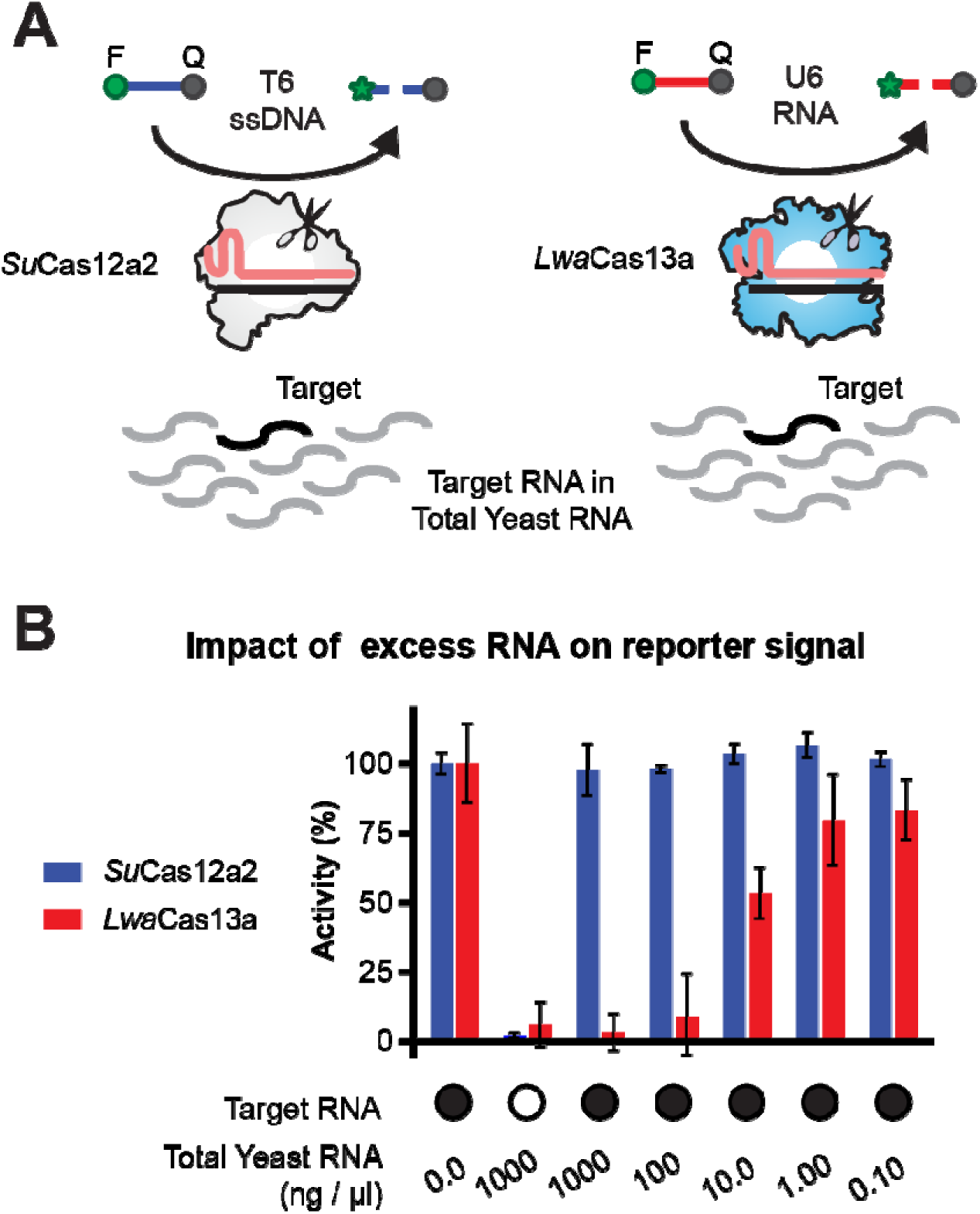
Cas12a2 collaterally cleaves fluorescent ssDNA probes in the presence of high concentrations of competing RNA. **A.** Cartoon schematic showing the comparison between Cas12a2 and Cas13a cleavage of their respective ssDNA or RNA probes in the presence of target RNA mixed with total yeast RNA. **B.** Competing total yeast RNA reduces Cas13a probe cleavage output after 30 minutes, while Cas12a2 cleavage is unaffected. Error bars represent one standard deviation of the mean.

After identifying the optimal ssDNA endpoint assay probe for Cas12a2, we compared the ability of Cas12a2 and Cas13a to cleave ssDNA T6 and ssRNA U6 reporter substrates, respectively, after recognizing a target RNA in the presence of competing total yeast RNA (**Figure 5A**). We found that the end-point fluorescence signal of Cas12a2 was consistent across all concentrations of total yeast RNA, while Cas13a activity was drastically reduced at the high yeast RNA concentrations of 1000 ng/μl and only returned to non-competition levels at yeast RNA concentrations of ∼0.1 ng/μl (**Figure 5B and Supplemental Table 5**). This result suggests that, in a diagnostic setup with a preamplification step or at high concentrations of non-reporter target RNA (∼nM-μM), competing RNAs may negatively affect the sensitivity of Cas13a-based diagnostics. In contrast, Cas12a2 is expected to retain its RNA diagnostic activity despite the presence of high levels of target or non-target RNA. Thus, Cas12a2’s strong preference for cleaving ssDNA substrates will enable DNA reporter-dependent RNA detection even in the presence of high concentrations of non-reporter RNAs. Indeed, work published while this manuscript was under revision used this property of Cas12a2 to develop an ultra-sensitive, amplification-free RNA detection tool (16).

## Discussion

Here, we present substrate binding and kinetic data that provide insight into how Cas12a2 selects and cleaves collateral substrates. RNA-activated Cas12a2 (ternary) binds ssRNA with the highest affinity of the three collateral substrate types, followed by ssDNA and dsDNA (**Figure 1**). Greater affinity for single-stranded substrates over dsDNA may be due to FAM-labeled single-stranded substrates binding to either the cleaved- or non-cleaved-strand binding sites of the open RuvC nuclease domain, while FAM-labeled dsDNA probes must locally unwind and bend to occupy both strand binding sites. Indeed, each binding site could have different affinities for ssRNA, ssDNA, and dsDNA, and it is possible that two labeled single-stranded probes could bind to sites where only a single dsDNA probe can fit. Understanding the affinities of each strand-binding site for all collateral substrates will be important for a full understanding of the collateral substrate cleavage mechanism and could help reveal the order of events associated with dsDNA bending, unwinding, and loading into the active site. Assays that can examine affinities at each binding site, such as single-molecule FRET, may reveal how differences in affinities at these sites influence collateral cleavage.

In our kinetic assays, ssRNA was cleaved with the highest turnover number (k_cat_) (**Table 2**). However, ssRNA cleavage exhibited lower catalytic efficiency than ssDNA cleavage, due to a fourfold higher Km (**Figure 1G-H and Table 2**). The higher k_cat_, yet lower catalytic efficiency for ssRNA cleavage compared to ssDNA cleavage, could be due to ssRNA associating and dissociating from the RuvC domain much faster than ssDNA, yet with a k_on_/k_off_ ratio indicating a smaller dissociation constant for ssRNA than for ssDNA and dsDNA. A faster k_off_ for ssRNA than for ssDNA would explain why PT-ssDNA is a better competitive inhibitor than PT-ssRNA. However, as the dissociation constants presented here were determined at equilibrium conditions, to examine this hypothesis, follow-on work is needed that determines the on and off rate constants of collateral substrates to activated Cas12a2.

Cas12a2 cleaves collateral ssDNA substrates with the greatest catalytic efficiency compared to collateral ssRNA and dsDNA. This activity is in line with the known biochemical activity of RuvC proteins, which resolve Holliday junctions by cleaving cross-over ssDNA (27–29), and displays similar turnover numbers and catalytic efficiencies to other single-subunit CRISPR-associated nucleases (11, 30). Additionally, the catalytic efficiencies for ssDNA and ssRNA that we determined are comparable to those recently published for Cas12a2 (10^5^ M^-1^ s^-1^) (16). However, the catalytic efficiency we determined for dsDNA cleavage is about 10-fold higher than previously reported. The large discrepancy in values is likely due to differences in the dsDNA probe design. We used a 10-nt probe with the same sequence as our single-stranded probes and placed the fluorophore and quencher on opposite sides of the probe (**Table 2**), whereas the previous work used a 30-nt probe with the fluorophore and quencher on the same side of the probe but on different strands (16). Since activated Cas12a2 binds only 11 base pairs at a time, and the fluorophore and quencher were positioned on the same side of the probe, it is likely that the 30-nt probe could be cleaved opposite the quencher and fluorophore without releasing fluorescence, thereby masking a larger turnover number and specificity constant for dsDNA cleavage.

Our kinetics work showed that dsDNA and ssDNA were cleaved with similar turnover numbers (**Table 2**). From this observation, we conclude that at enzyme-saturating concentrations, the fluorophore-labeled strand of the dsDNA probe is cleaved just as fast as FAM-labeled ssDNA. Importantly, this similarity indicates that the FAM label was removed from the dsDNA probe, about as fast as from ssDNA, regardless of whether the labeled strand was initially bound to the cleaved or non-cleaved strand-binding site. We propose that these results indicate a two-strand cleavage mechanism in which both strands can sample the RuvC cleavage site after initial unwinding and bending of the duplex (**Figure 6**). This model is supported by plasmid cleavage data showing production of double-stranded breaks, or linear bands, within the first minute.

**Figure 6.**
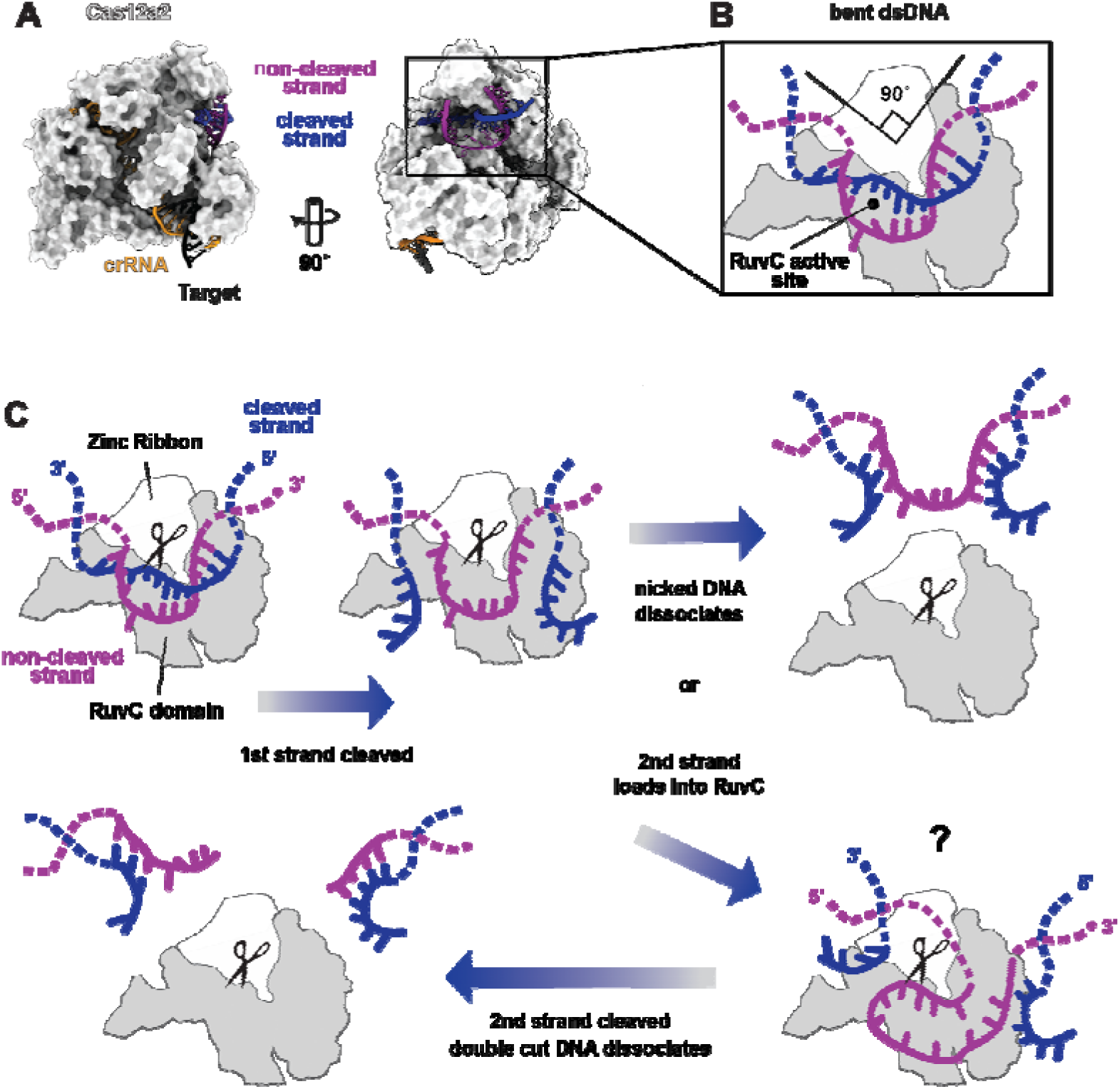
Model of collateral dsDNA cleavage mechanism. (A) Quaternary structure of Cas12a2 bound to an 11-nt collateral Phosphorothioated dsDNA substrate (PDB 8D4A). (B) 2D diagram of dsDNA bound by the Zn ribbon and RuvC nuclease domain. The 90° bend in the dsDNA is indicated, as well as the location of the RuvC nuclease active site. (C) Model depicting a mechanism of collateral dsDNA cleavage. Our kinetic and plasmid cleavage data suggest that after the first strand is cleaved, the nicked DNA dissociates from Cas12a2 or the second strand of the duplex loads into the RuvC active site for cleavage with an unknown mechanism indicated with a question mark. The depiction of how the second strand may load into the RuvC active site was inspired by recent structures demonstrating how Cas12a1 cleaves the target strand of bound dsDNA (31).

However, the cleavage products also indicate wild-type Cas12a2 cuts both strands of supercoiled DNA only half the time (50% open circle and 50% linear) (**Figure 3**) suggesting a mechanism where after one strand of the dsDNA substrate is cleaved, there is an equal probability that second strand docks into the cleavage site to be cleaved, or Cas12a2 dissociates from the nicked DNA (**Figure 6**). Although this mechanism requires further investigation, the highly bent duplex DNA bound into the active site of the quaternary Cas12a2 structure positions the non-cleaved strand of the DNA duplex near the active site, suggesting the structural rearrangements that occur upon cleavage of the first strand may allow the second strand to quickly bind into the active site. Loading of the second strand into the RuvC active site may share mechanistic similarities with the target-strand loading and cleavage mechanism of Cas12a1 (31).

Our cleavage analysis with aromatic clamp mutants suggests that the aromatic clamps may play a role in the proposed two-strand cleavage mechanism. Mutating aromatic residue Y1069 that clamps the cleaved strand, or Y465 that clamps the non-cleaved strand, impaired collateral cleavage of our dsDNA probe, but not ssRNA or ssDNA (**Figure 3A-E**). Additionally, in our plasmid cleavage assay, mutating either of the aromatic residues that clamp the cleaved strand, F1092 or Y1069, converted Cas12a2 into a nickase, similar to Cas12a1 (13) (**Figure 3F-I**). In contrast, mutating the residues that interact with the non-cleaved strand either had no observable effect on cleavage (Y1080) or impaired the cleavage rate, but did not change the type of cleavage products formed (Y465) (**Figure 3J**). Combined, these kinetic and plasmid cleavage data suggest that the cleaved-strand aromatic clamp is essential for cleavage of the second DNA strand, perhaps through binding the second strand into the active site before product dissociation.

Phosphorothioated PT-ssRNA and PT-ssDNA substrates competitively inhibited cleavage of single-stranded substrates, and inhibited dsDNA cleavage with mixed inhibition (**Figure 4F-G and Table 3**). Mixed inhibition is observed when an inhibitor binds to the unbound enzyme with an inhibition constant K_i_ and the substrate-bound complex with a different inhibition constant K_i_′. Both PT-ssRNA and PT-ssDNA inhibited dsDNA cleavage with smaller K_i_ values than for ssDNA cleavage, and PT-ssDNA inhibited dsDNA cleavage with a 2-fold smaller K_i_ than PT-ssRNA, suggesting dsDNA cleavage is inhibited more strongly than single-stranded substrate cleavage, and PT-ssDNA is the stronger inhibitor. The smaller K_i_ values (stronger inhibition) for dsDNA cleavage likely reflect single-stranded PT inhibitors that preclude dsDNA cleavage when bound to either the cleaved or non-cleaved strand binding sites in the RuvC nuclease domain, whereas ssDNA or ssRNA cleavage is inhibited only when the inhibitors bind the cleaved strand binding site. In addition to determining K_i_ values, our fit to a mixed inhibition model revealed PT-ssRNA and PT-ssDNA inhibit dsDNA cleavage with comparable K_i_′ values, yet with PT-ssRNA showing a slightly higher K_i_′, or slightly less inhibition. Since our inhibitors are the exact same nucleotide sequence as our probes, a single-stranded PT-ssRNA or PT-ssDNA could readily form a triple-helix structure via Hoogsteen base-pairing with the FAM-labeled dsDNA probe, or could form Watson-Crick base pairs with melted bases, likely affecting the flexibility of bound dsDNA probes and their ability to fit into the Cas12a2 binding pocket. Therefore, these data suggest that, while most of the PT-ssDNA or PT-ssRNA inhibitors likely bind competitively to the RuvC domain, a smaller fraction may bind the quaternary Cas12a2:crRNA:target:dsDNA complex (ternary Cas12a2 – dsDNA probe complex), thereby impairing cleavage of bound dsDNA. Future work with inhibitor sequences that are less likely to base-pair with dsDNA will allow us to better understand how single-stranded substrates compete with dsDNA substrates for Cas12a2 collateral cleavage.

Here, we demonstrated that target RNA-triggered Cas12a2 effectively cleaves collateral ssDNA substrates, even in the presence of high concentrations of background non-target RNA. Notably, Cas13a-based RNA targeting that relies on collateral cleavage of an RNA probe was impaired by competing RNA, suggesting that Cas12a2 may be a more optimal enzyme for diagnostic applications in the presence of high concentrations of background RNA. We hypothesize that this phenomenon stems from ssRNA interacting with the RuvC domain, with faster k_on_ and k_off_ values than DNA, due to possible differences in how RNA substrates bind at the active site. However, additional biochemical studies and molecular structures of Cas12a2 bound to collateral RNA are needed to fully examine this hypothesis. Collectively, our data provide insight into the cleavage preferences and mechanisms of Cas12a2’s collateral nuclease activity, which are essential for developing Cas12a2-based applications.

## Data availability

Raw Illumina sequencing reads have been deposited at NCBI Sequence Read Archive (SRA) under submission BioProject PRJNA1203263 and are publicly available. Any additional raw data associated with this paper are available from the lead author upon request.

## Supplementary data

Supplemental Figures S1 - S10 and Tables 1 - 5 are found in the Supplementary Information File.

## Author Contributions

T.H., S.K., and R.N.J. conceived the experiments. T.H., S.K., and B.N. expressed and purified Cas12a2 and mutants. T.H. and S.K. collected fluorescence anisotropy data. T.H., S.K., and S.M. performed kinetic experiments. T.H., D.K., and A.T. collected the plasmid cleavage sequencing data. T.H. and S.K. performed competition experiments. S.K. performed plasmid cleavage assays. S.M. screened diagnostics probes, and T.H. compared total yeast RNA competition with Cas12a2 and Cas13a. T.H. wrote the initial draft of the manuscript. S.K. and R.N.J. wrote the revised manuscript. T.H., S.K., S.M., C.L.B., and R.N.J. provided critical feedback on the final manuscript. R.N.J. and C.L.B. secured funding for the research.

## Acknowledgements

We thank Dr. Nicholas E. Dickensen of Utah State University for generously providing access to and technical support with the Synergy H4 Hybrid plate reader used in this study. We also thank Drs. Nicholas E. Dickenson, Sean J. Johnson, and Lance C. Seefeldt of the Department of Chemistry and Biochemistry at Utah State University for their review and comments on the manuscript before submission. We would also like to thank the reviewers of the first draft of this manuscript, who provided critical advice and guidance that substantially improved this work.

Support for this work was provided by the National Institutes of Health, the National Institute of General Medical Sciences [R35GM138080] (RNJ); the European Research Council Proof-of-Concept award [101158249] (C.L.B.); and Helmholtz Center for Infection Research (HZI) Singh-Chhatwal-Postdoctoral Fellowship (S.M.). The contents of this publication are solely the responsibility of the authors and do not necessarily represent the official views of the NIH, NIGMS, or ERC.

