## Supplemental Data for "Target RNA-triggered CRISPR-Cas12a2 Preferentially Cleaves Collateral DNA over RNA"

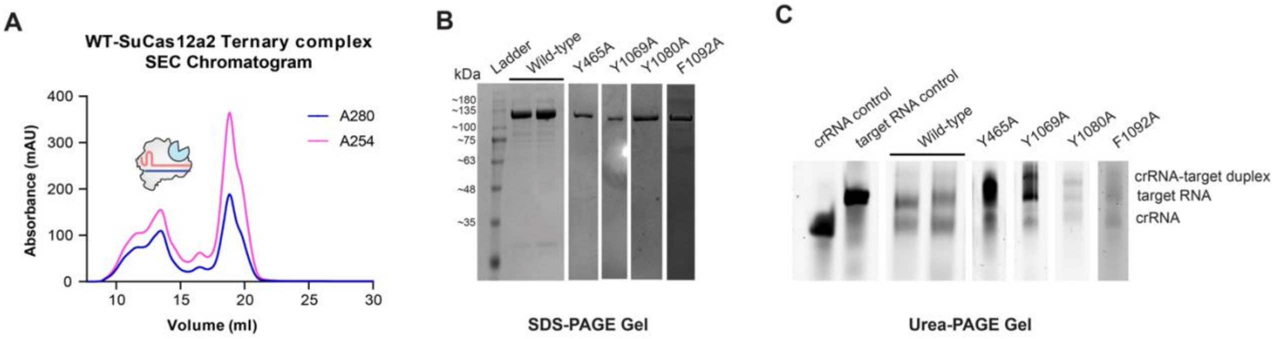


**Supplemental Figure S1. Purification of ternary complexes of wild-type and its aromatic clamp mutants. A.** An example of a size exclusion chromatogram for the purification of the ternary complex of wild-type SuCas12a2. **B.** SDS-PAGE gel confirms the presence of purified wild-type and mutant Cas12a2 proteins (Y465A, Y1069A, Y1080A, and F1092A). **C.** Urea-PAGE gel confirms the presence of crRNA and target RNA. Control lanes show crRNA alone and target RNA alone. Together, these gels confirm the formation of ternary complexes.

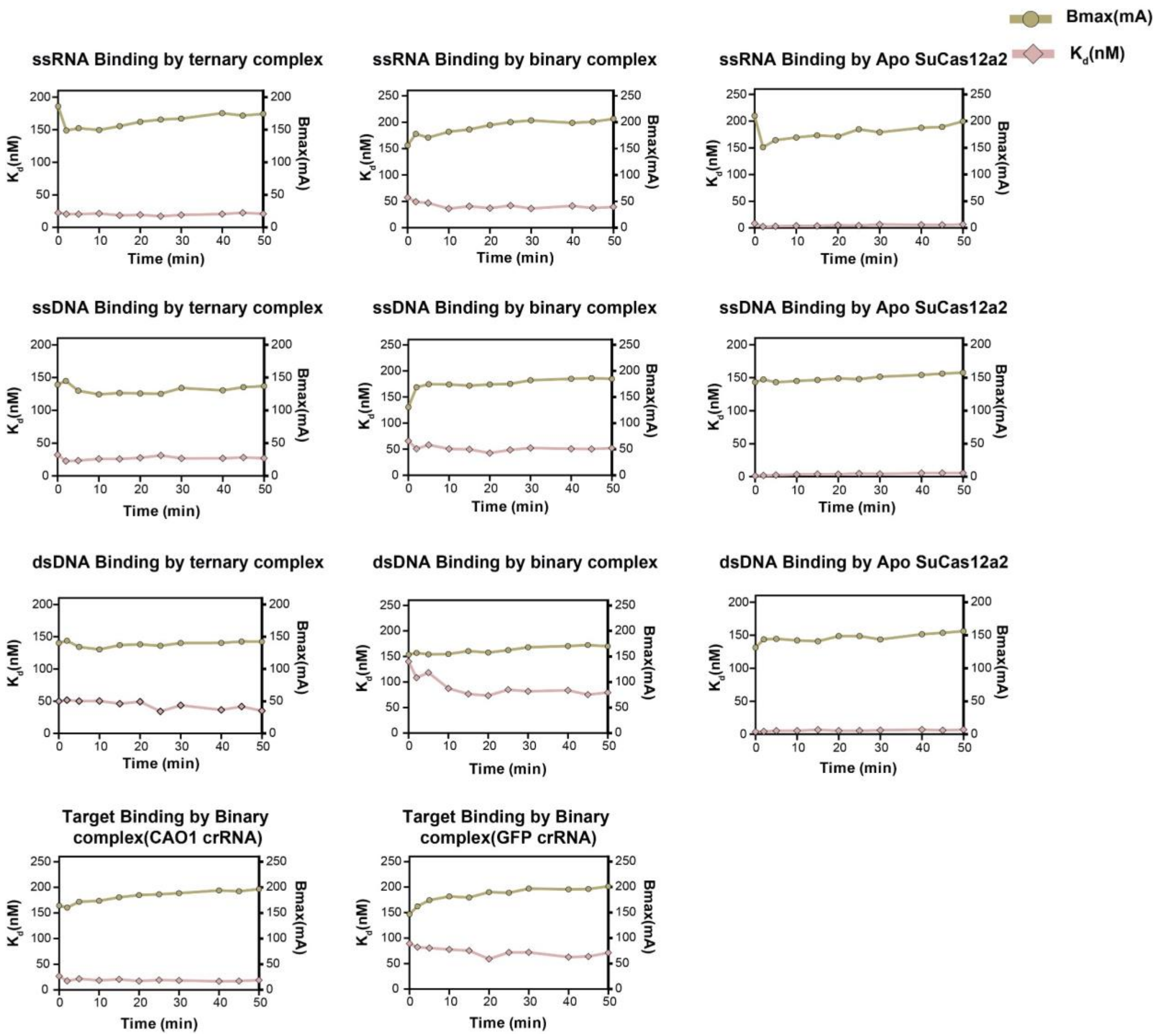


**Supplemental Figure S2. Time-dependent binding analysis of collateral nucleic acid substrates and target RNA.** Curves showing the dissociation constant ($K_d$) and $B_{max}$ values over time, indicating the time it takes to reach equilibrium binding of different collateral nucleic acid substrates with the apo, binary, and ternary complexes. The bottom panel shows target RNA binding by two distinct binary complexes assembled with different guides (CAO1 and GFP).

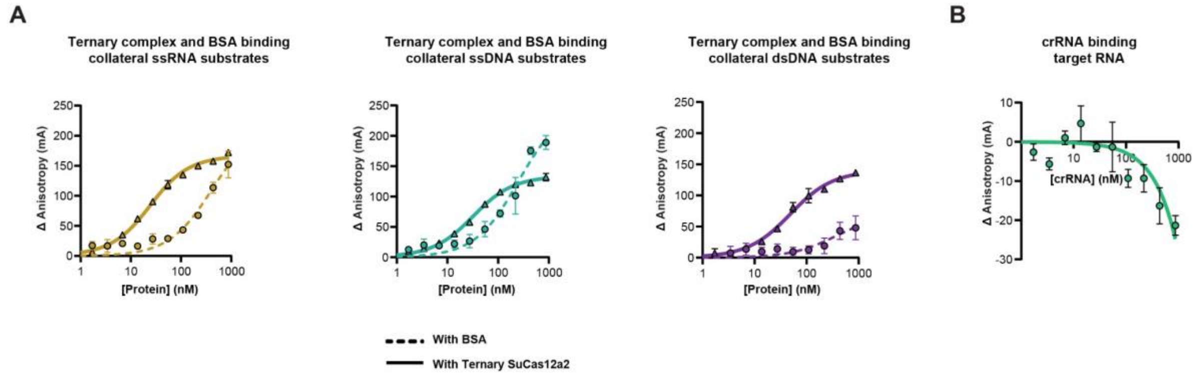


C

| Binding parameters determined by fluorescence anisotropy | | |
|---|---|---|
| **Substrate** | **Bmax (mA)** | **Apparent $K_d$ (nM)** |
| **BSA** | | |
| Collateral ssRNA | 220 ± 20 | 410 ± 90 |
| Collateral ssDNA | 250 ± 20 | 240 ± 40 |
| Collateral dsDNA | 70 ± 40 | 350 ± 560 |
| **crRNA** | | |
| Target RNA | N/A | N/A |
| Nucleic acid substrates used are non-hydrolyzable, phosphorothioated (PT) substrates. | | |

**Supplemental Figure S3. Fluorescence anisotropy data comparison for Cas12a2 and BSA collateral substrate binding. A**. Binding curves of different collateral substrates by the ternary complex of SuCas12a2 (solid lines) and BSA (dashed lines) are shown. These curves were used to calculate collateral substrate dissociation constants. **B.** The binding curve of FAM-labeled target RNA to crRNA without Cas12a2 is shown. **C.** Table representing $B_{max}$ and $K_d$ values for collateral substrates binding by BSA and target RNA binding by crRNA.

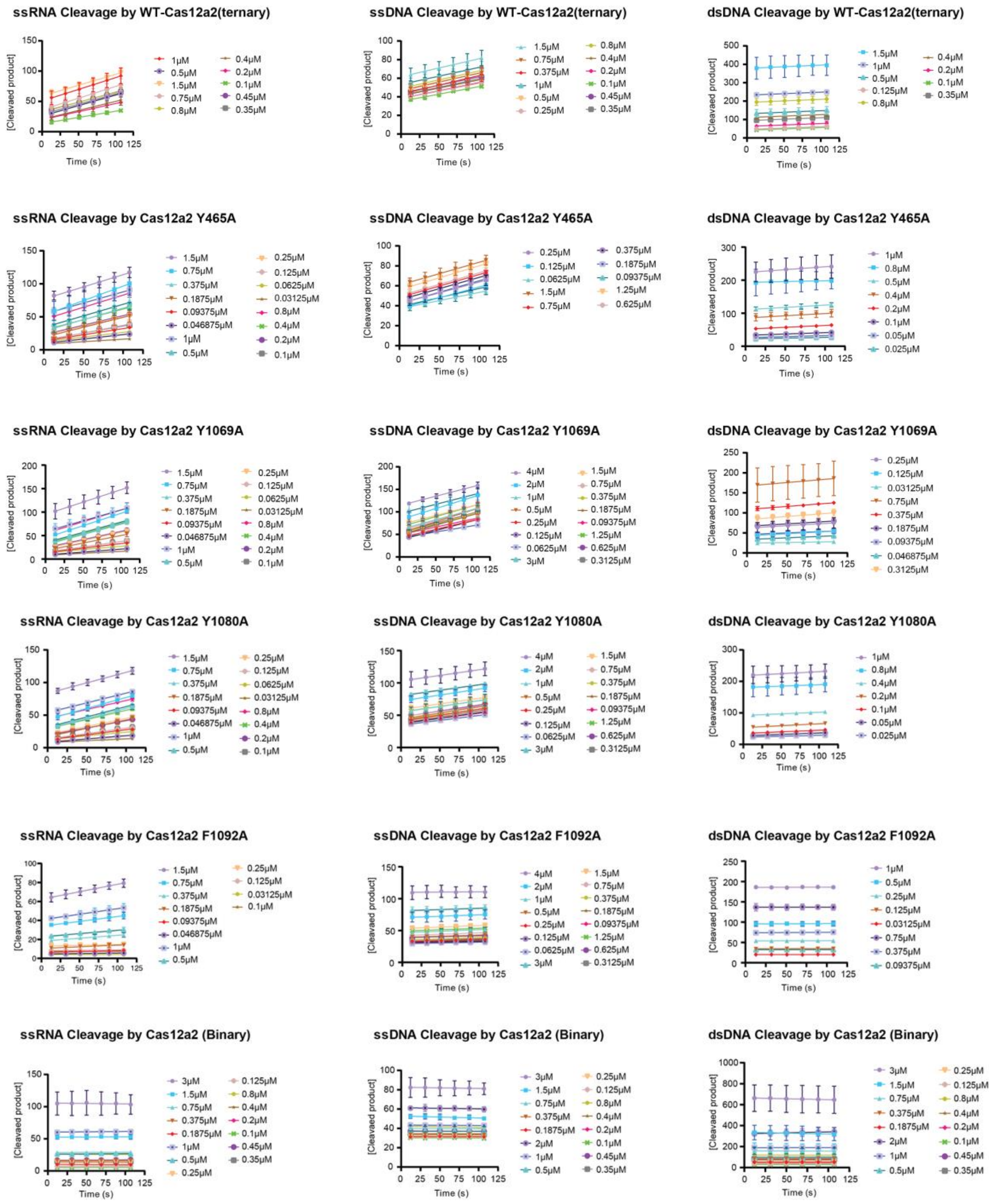


**Supplemental Figure S4-1. Progress curves for collateral substrate cleavage by WT Cas12a2 and aromatic clamp mutants.** Linear progress curves showing cleavage of collateral substrates by wild-type (WT) Cas12a2 and Cas12a2 aromatic clamp mutants over 5 time points at intervals of 19 seconds of reaction. The left panel shows ssRNA probe cleavage at varying ssRNA probe concentrations. The middle panel shows ssDNA probe cleavage at varying ssDNA probe concentrations. The right panel shows dsDNA probe cleavage at varying dsDNA probe concentrations. Progress curves for higher dsDNA probe concentrations (> 1.5 μM) were not included due to inconsistencies in the fluorescence signal. Error bars represent the standard deviation of the mean from three independent replicates.

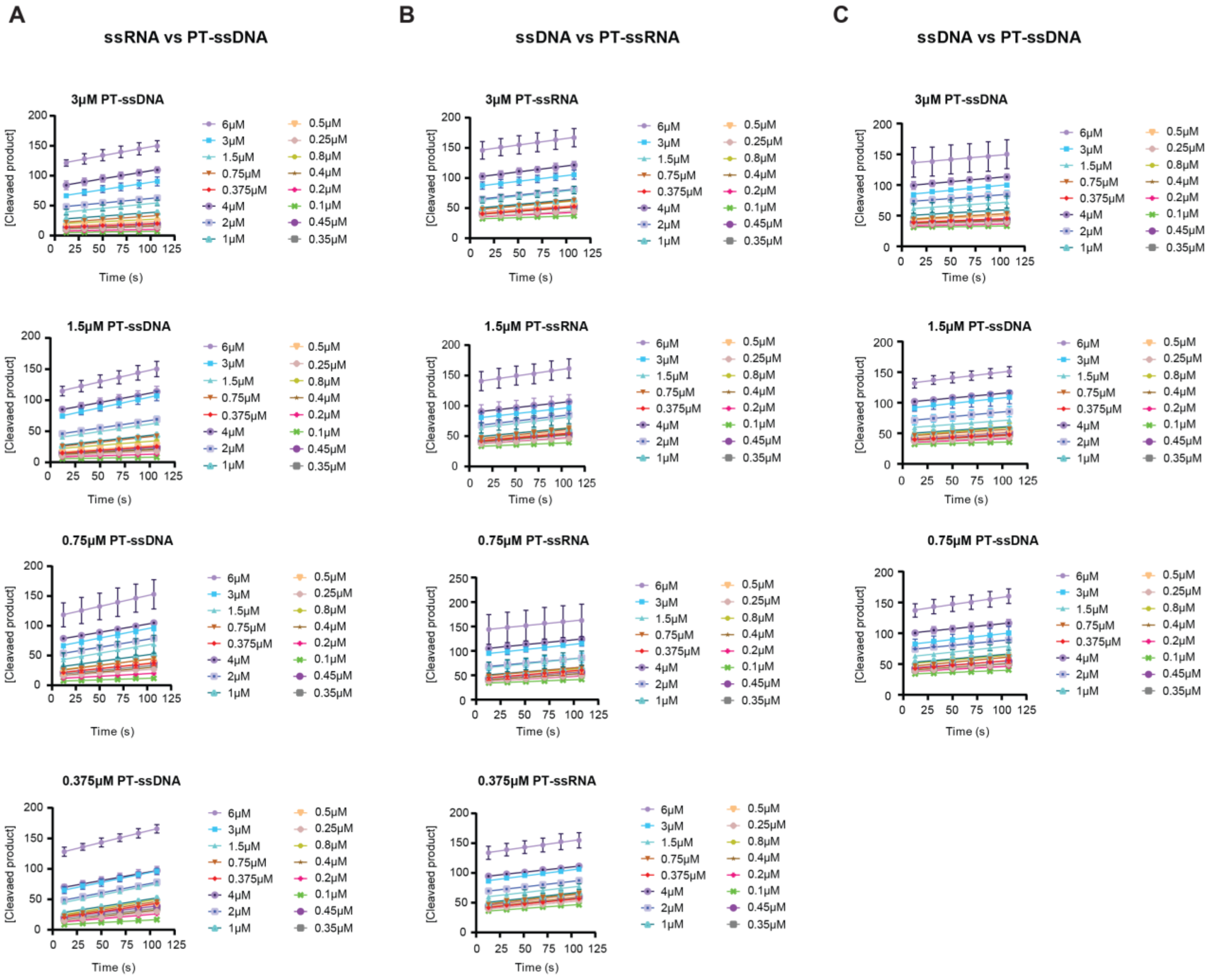

D

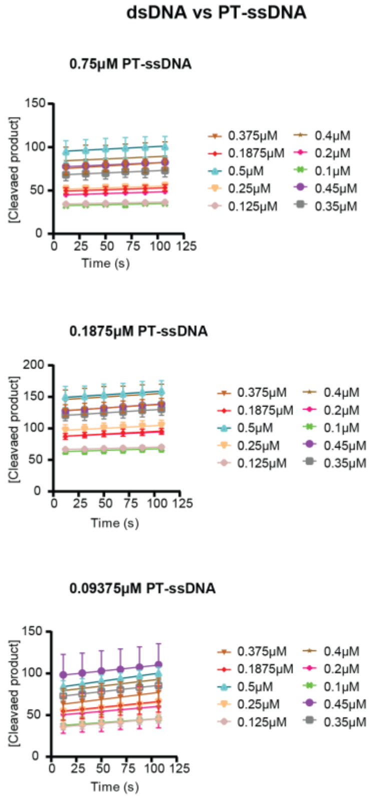


E

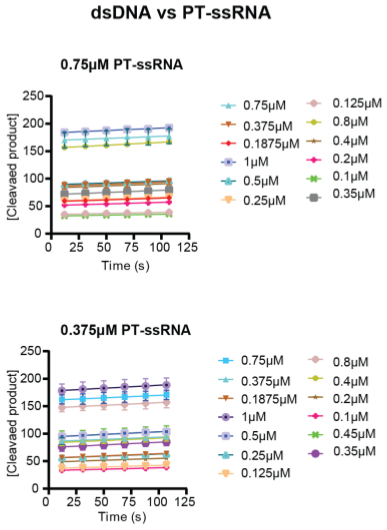


**Supplemental Figure S4-2. Progress curves for collateral substrate cleavage by WT Cas12a2 in the presence of different concentrations of inhibitors.** Linear progress curves showing cleavage of collateral substrates by wild-type (WT) Cas12a2 in the presence of phosphorothioate inhibitors over the initial 5 time points at an interval of 19 seconds of reaction. **Panel A** shows ssRNA cleavage in the presence of various concentrations of PT-ssDNA. **Panel B** shows ssDNA cleavage in the presence of various concentrations of PT-ssRNA. **Panel C** shows ssDNA cleavage in the presence of various concentrations of PT-ssDNA. **Panel D** shows dsDNA cleavage in the presence of various concentrations of PT-ssDNA. **Panel E** shows dsDNA cleavage in the presence of various concentrations of PT-ssRNA. Progress curves for higher dsDNA concentrations (> 1.5 μM) were not included due to inconsistencies in the fluorescence signal. Error bars represent the standard deviation of the mean from independent replicates.

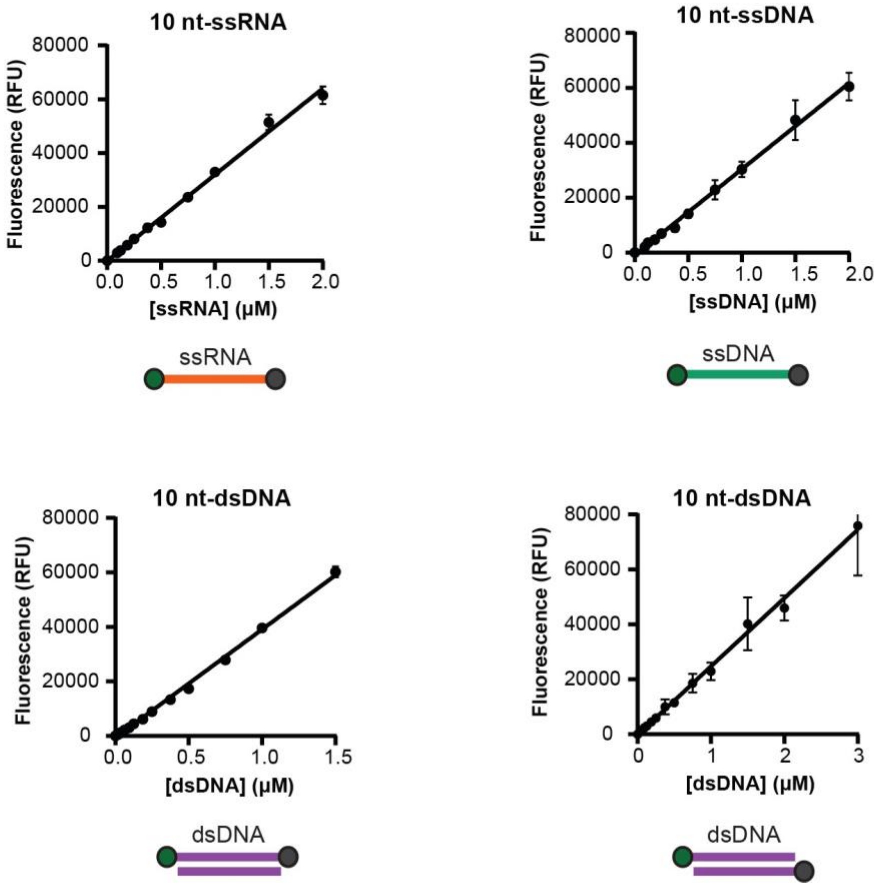


**Supplemental Figure S5. Standard curves for different fluorescent reporter substrate quantification**. The standard curves for various single-stranded RNA (ssRNA), single-stranded DNA (ssDNA), and double-stranded DNA (dsDNA) were generated by plotting fluorescence intensity against substrate concentration. The fluorescence signal observed in the kinetic assays was converted to a molar concentration using a linear fit of these standard curves.

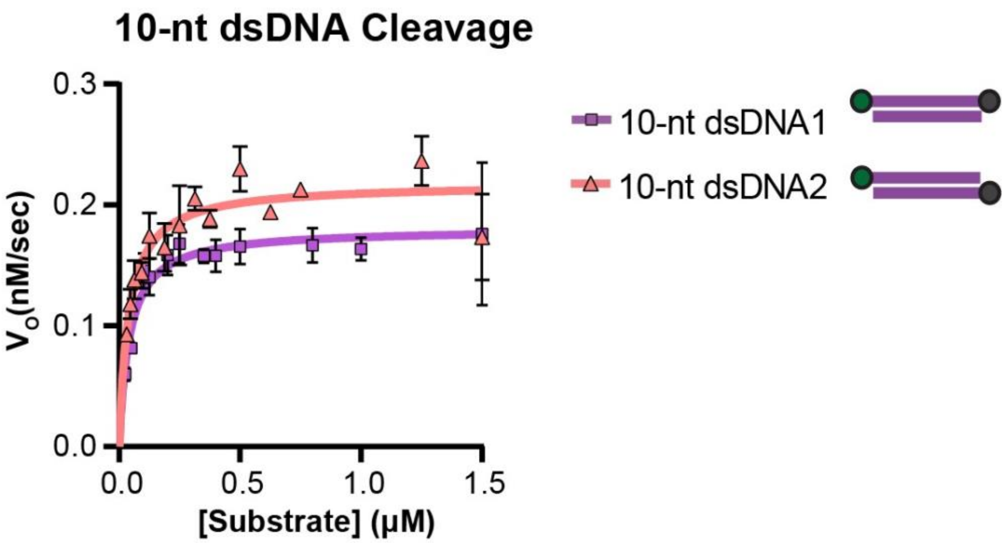


**Supplemental Figure S6. Comparison of dsDNA substrate cleavage kinetics fitted to the Michaelis-Menten model.** Cleavage kinetics of different dsDNA substrates were analyzed and fitted to the Michaelis-Menten equation. The pink curve represents cleavage of dsDNA substrates containing the fluorophore and quencher on opposite strands. The purple curve represents cleavage of dsDNA substrates containing the fluorophore and quencher on the same strand. Data points were fit to the Michaelis-Menten model using nonlinear regression.

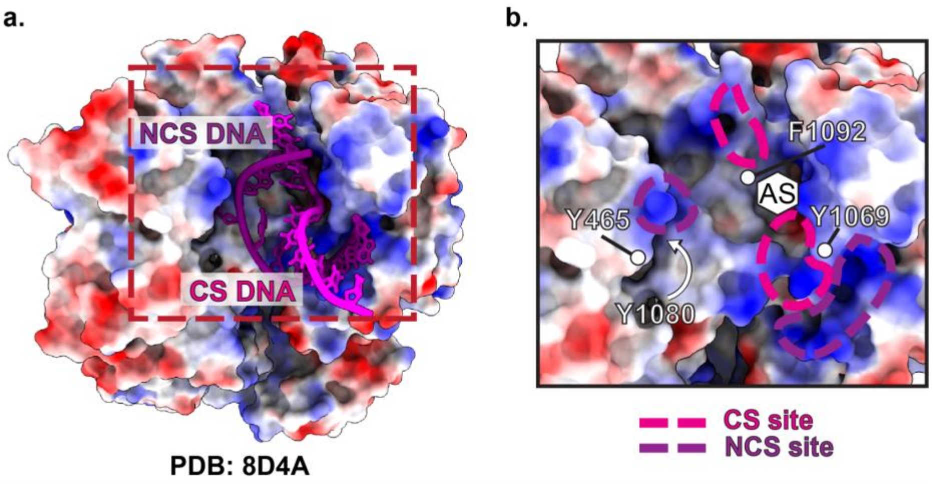


**Supplemental Figure S7. Structural overview of Cas12a2 bound to cleaved (CS) and non-cleaved (NCS) strands of double-stranded DNA. a.** The electrostatic surface representation (blue positive, red negative) of Cas12a2 (PDB: 8D4A) bound to both cleaved (CS, magenta) and non-cleaved (NCS, purple) DNA strands. The cleaved and non-cleaved strands are positioned within the binding pocket of the protein. **b,** Close-up view highlighting key amino acid residues involved in binding the cleaved and non-cleaved DNA strands. Each strand is coordinated by a pair of aromatic clamps (Y1069 and F1092 for CS, and Y465 and Y1080 for NCS). The active site (AS) is marked, and dashed lines indicate binding regions for the CS site (magenta) and NCS site (purple).

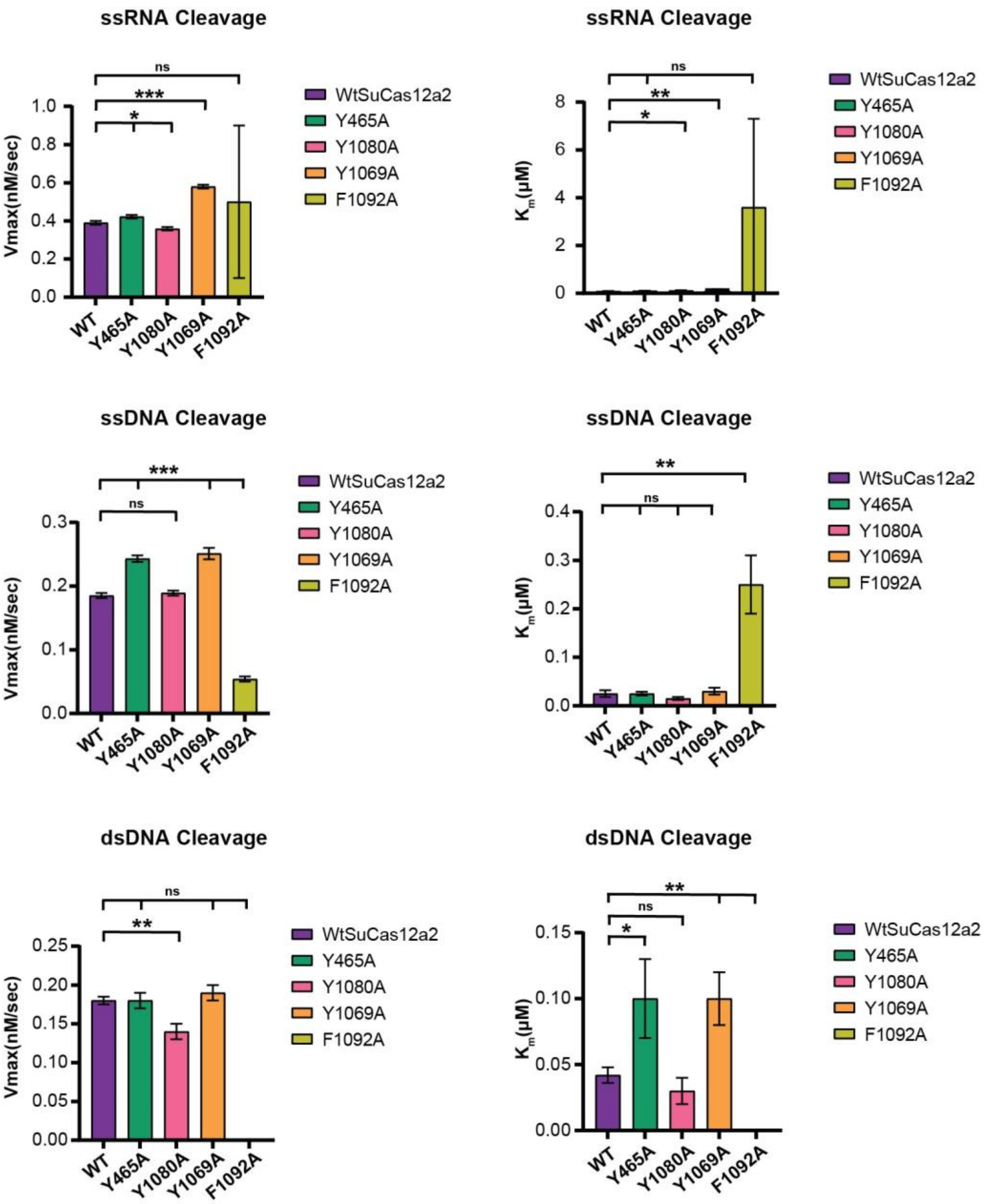


**Supplemental Figure S8. Comparison of $V_{max}$ and $K_m$ values for different nucleic acid substrates cleavage by wild-type Cas12a2 and its aromatic clamp mutants.** Bar graphs show changes in $V_{max}$ (left panels) and Km (right panels) across different mutants, with each mutant represented by a different color. Error bars represent the standard deviation of the mean, with statistical significance denoted as *: $p < 0.05$, **: $p < 0.01$, ***: $p<0.001$, ****: $p<0.0001$ with a student t-test.

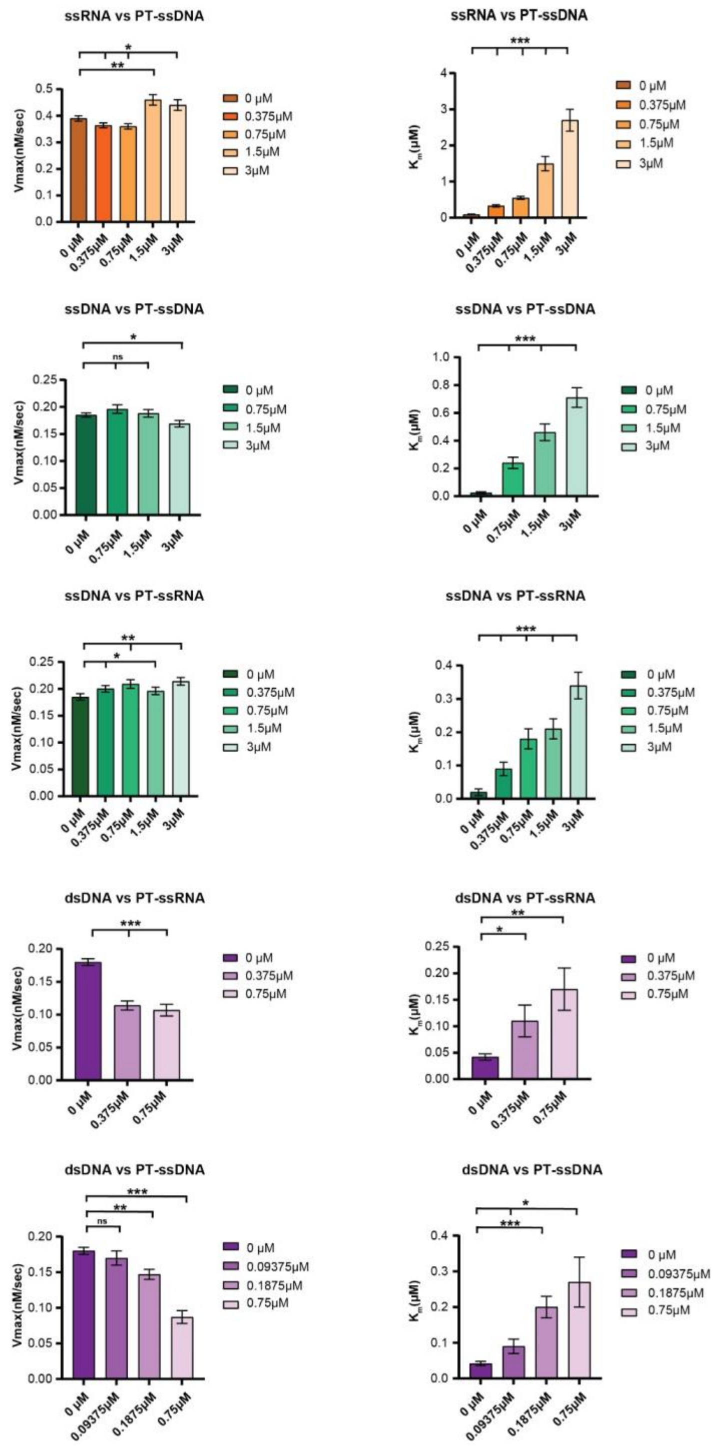


**Supplemental Figure S9. Effects of non-hydrolyzable inhibitor concentrations on $K_m$ and $V_{max}$ values of different nucleic acid cleavage by wild-type Cas12a2.** Bar graphs show changes in $V_{max}$ (left panels) and $K_m$ (right panels) in the presence of increasing concentrations of inhibitor, represented by decreasing shades of respective color. Error bars represent the standard deviation of the mean, with statistical significance denoted as *: $p < 0.05$, **: $p < 0.01$, ***: $p<0.001$, ****: $p<0.0001$ with a student t-test.

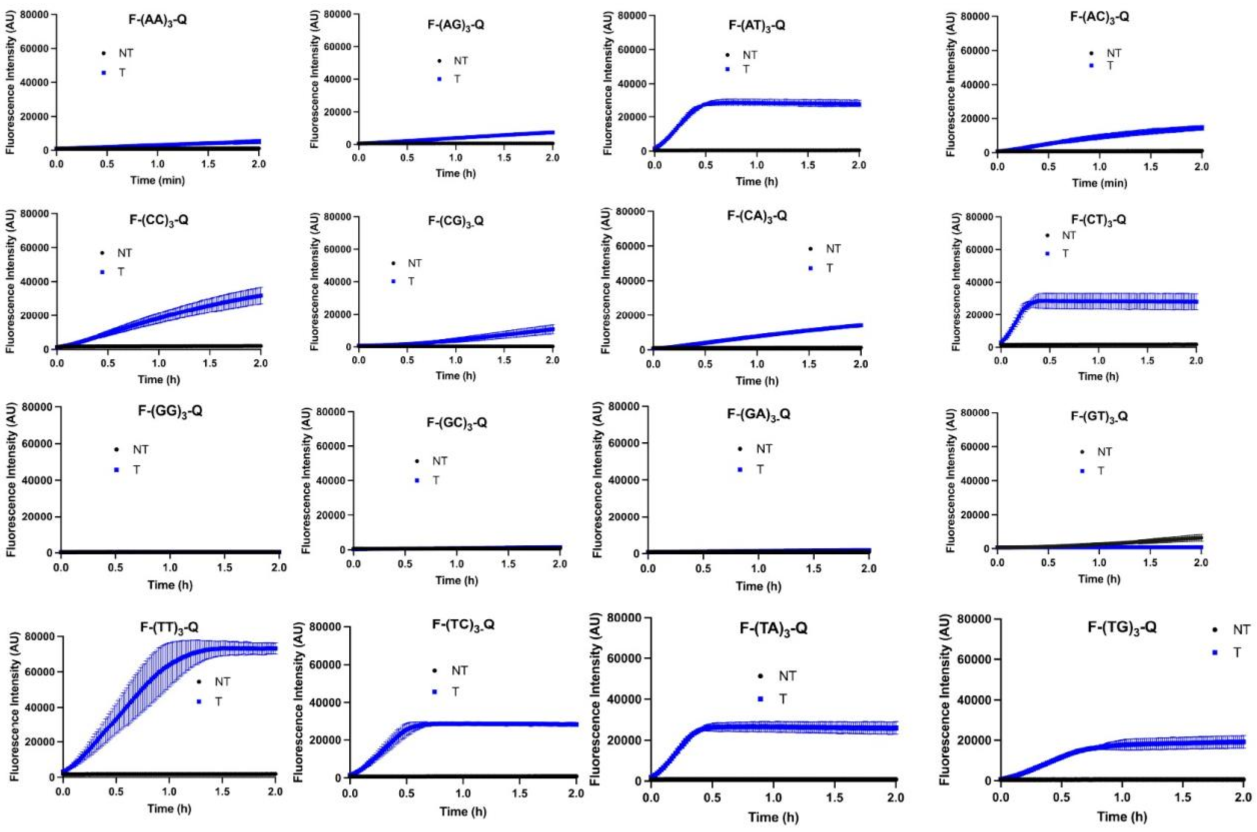


**Supplemental Figure S10. Optimization of ssDNA reporter probe length and sequence for RNA detection by Cas12a2.** Screening of 6-nt mono- and dinucleotide sequence combinations to identify the optimal reporter sequence for Cas12a2-mediated RNA detection. Error bars represent the standard deviation of the mean.

**Supplemental Table 1.** Sequences of Substrates and Primers

Substrates

| **Substrates** | **Sequences** |
|---|---|
| 43 nt- CAO1 crRNA | 5'- AAUUUCUACUAUUGUAGAUUGGAGCAACACCUGAAGGAAGGCU – 3' |
| 42 nt- GFP crRNA | 5'- UAAAUUUCUACUAUUGUAGAUCGUCGCCGUCCAGCUCGACCA -3' |
| 67 nt- target RNA | 5'- GCUGCCGCACUUGCUCAUCAAGCCUUCCUUCAGGUGUUGCUCCAGAAAGGUGAGUUCUUCUUGUUGU -3' |

Substrates for Binding assay

| **Substrates** | **Sequences (With 5'- Fluorescein )** |
|---|---|
| 53 nt- PT- target RNA | 5'-**F**-rC*rG*rC*rA*rC*rU*rU*rG*rC*rU*rC*rA*rU*rC*rA*rA*rG*rC*rC*rU*rU*rC*rC*rU*rU*rC*rA*rG*rG*<br>rU*rG*rU*rU*rG*rC*rU*rC*rC*rA*rG*rA*rA*rA*rG*rG*rU*rG*rA*rG*rU*rU*rC*rU-3' |
| 53 nt-PT-ssRNA | 5'-**F**-rC*rA*rG*rA*rG*rA*rU*rA*rA*rG*rU*rG*rA*rC*rG*rC*rG*rC*rG*rG*rC*rG*rA*rG*rU*rG*rG*rC*rG*<br>rC*rG*rC*rC*rA*rC*rG*rU*rC*rG*rG*rA*rA*rA*rU*rC*rU*rA*rG*rA*rG*rG*rC*rG-3' |
| 51 nt-PT-ssDNA | 5'-**F**-A*A*C*T*G*A*T*A*T*G*A*C*A*A*T*T*G*C*G*C*G*T*A*G*C*A*C*G*A*C*G*A*C*G*A*T*A*T*G*<br>A*C*A*C*T*T*G*C*G*C*A*T-3' |
| 51 bp-PT-dsDNA | 5'-**F**-A*A*C*T*G*A*T*A*T*G*A*C*A*A*T*T*G*C*G*C*G*T*A*G*C*A*C*G*A*C*G*A*C*G*A*T*A*T*G*<br>A*C*A*C*T*T*G*C*G*C*A*T-3'<br>3'-T*T*G*A*C*T*A*T*A*C*T*G*T*T*A*A*C*G*C*G*C*A*T*C*G*T*G*C*T*G*C*T*G*C*T*A*T*A*C*T*<br>G*T*G*A*A*C*G*C*G*T*A-5' |

Substrates for Kinetic assay

| **Substrates** | **Sequences ( With 5'- Fluorescein and 3'- Iowa Black Quencher)** |
|---|---|
| 10 nt-ssRNA | 5'-**F**-UGUAACGACC-**Q**-3' |
| 10nt- ssDNA | 5'-**F**-TGTAACGACC-**Q**-3' |
| 10 bp-dsDNA | 5'-**F**-TGTAACGACC-**Q**-3'<br>3'- ACATTGCTGG -5' |

| **Primers** | **Sequences** |
|---|---|
| **Adapter Sequences** | |
| i5 | biotin-ACACTCTTTCCCTACACGACGCTCTTCCGATCT |
| i5-R | GATCGGAAGAGCGTCGTGTAGGGAAAGAGTGT |
| i7 | GTGACTGGAGTTCAGACGTGTGCTCTTCCGATCT |
| i7-R | phos-GATCGGAAGAGCACACGTCTGAACTCCAGTCAC |
| **Indexing Primers** | |
| i5 | AATGATACGGCGACCACCGAGATCTACACnnnnnnnnACACTCTTTCCCTACACGACG |
| i7 | CAAGCAGAAGACGGCATACGAGATnnnnnnnnGTGACTGGAGTTCAGACGTGTG |

Inhibitors for Competition assay

| **Inhibitors** | **Sequences** |
|---|---|
| 10 nt-PT-ssRNA | 5'-rU*rG*rU*rA*rA*rC*rG*rA*rC*rC-3' |
| 10 nt-PT-ssDNA | 5'-T*G*T*A*A*C*G*A*C*C-3' |

**F represents 5'- Fluorescein, Q represents 3'- Iowa Black Quencher, * represents phoshphorothioate bond**

**Supplemental Table 2.** Representative Cas12a2 cut-site sequence against pUC19.

| Position | Sequence* | Frequency |
|---|---|---|
| 1302 | aaaaagg.aagagta | 562 |
| 2137 | atcgctg.agatagg | 303 |
| 944 | gtacaat.ctgctct | 196 |
| 1602 | tcagaat.gacttgg | 190 |
| 1604 | agaatga.cttggtt | 183 |
| 418 | aggaagc.ggaagag | 158 |
| 2580 | ggctgct.gccagtg | 155 |
| 426 | gaagagc.gcccaat | 146 |
| 637 | gaaacag.ctatgac | 136 |
| 1912 | gcttccc.ggcaaca | 135 |
| 425 | ggaagag.cgcccaa | 132 |
| 1291 | caataat.attgaaa | 132 |
| 298 | ttgctca.catgttc | 129 |
| 1709 | cggccaa.cttactt | 126 |
| 946 | acaatct.gctctga | 120 |
| 638 | aaacagc.tatgacc | 118 |
| 1686 | cataacc.atgagtg | 111 |
| 871 | gaatggc.gcctgat | 106 |
| 1299 | ttgaaaa.aggaaga | 105 |
| 1313 | gtatgag.tattcaa | 105 |
| 1075 | atgtgtc.agaggtt | 104 |
| 579 | tttatgc.ttccggc | 102 |
| 1896 | cgaacta.cttactc | 102 |
| 1236 | ctaaata.cattcaa | 101 |
| 1873 | acgttgc.gcaaact | 101 |

*: The position of cleavage is indicated by a "." .

**Supplemental Table 3.** Standard Curve Equations for Fluorescent Reporter Substrate Quantification

| Substrate | Equation |
|---|---|
| **10 nt-ssRNA** | y = 32050x – 72.4 |
| **10 nt-ssDNA** | y = 31293x – 893.2 |
| **10 bp-dsDNA ( with 5'-FAM and 3'-Quencher on same strand)** | y = 39714x – 642.1 |
| **10 bp-dsDNA ( with 5'-FAM and 5'-Quencher on different strand)** | y = 24872x – 163.7 |
| where y is fluorescence (RFU); x is concentration in µM. | |

**Supplemental Table 4. Summary of Experimentally Measured Kinetic Parameters collected from different sources**

| Enzyme | target | Reporters (5'-3') | $k_{cat}$ ($s^{-1}$) | $K_m$(nM) | $k_{cat}/K_m$ ($M^{-1}$ $s^{-1}$) | References |
|---|---|---|---|---|---|---|
| LbCas12a-1 | ssDNA | | 0.58 | 310 | $1.9 \times 10^6$ | |
| LbCas12a-1 | dsDNA | | 0.38 | 370 | $1.0 \times 10^6$ | |
| LbCas12a-2 | ssDNA | | 0.048 | 230 | $2.1 \times 10^5$ | |
| LbCas12a-2 | dsDNA | | 0.28 | 680 | $4.2 \times 10^5$ | |
| LbCas12a-3 | ssDNA | | 0.29 | 240 | $1.2 \times 10^6$ | |
| LbCas12a-3 | dsDNA | | 0.19 | 320 | $5.9 \times 10^5$ | |
| LbCas12a-4 | ssDNA | | 0.27 | 180 | $1.4 \times 10^6$ | |
| LbCas12a-4 | dsDNA | ssDNA | 0.32 | 260 | $1.2 \times 10^6$ | |
| LbCas12a-5 | ssDNA | F-TTATTATT-Q | 0.098 | 120 | $8.5 \times 10^5$ | |
| LbCas12a-5 | dsDNA | | 0.078 | 82 | $9.5 \times 10^5$ | |
| LbCas12a-6 | ssDNA | | 0.3 | 260 | $1.2 \times 10^6$ | |
| LbCas12a-6 | dsDNA | | 0.56 | 490 | $1.1 \times 10^6$ | **Huyke et al,2022** |
| LbCas12a-7 | ssDNA | | 0.022 | 450 | $4.9 \times 10^4$ | |
| LbCas12a-7 | dsDNA | | 0.028 | 870 | $3.3 \times 10^4$ | |
| AsCas12a-1 | dsDNA | | 0.46 | 280 | $1.6 \times 10^6$ | |
| AsCas12a-2 | ssDNA | | 1.3 | 420 | $3.0 \times 10^6$ | |
| AsCas12a-2 | dsDNA | | 0.9 | 1100 | $8.2 \times 10^5$ | |
| AapCas12b-1 | ssDNA | | 0.13 | 660 | $2.0 \times 10^5$ | |
| AapCas12b-1 | dsDNA | | 0.049 | 920 | $5.3 \times 10^4$ | |
| LwaCas13a-1 | RNA | | 2.4 | 2100 | $1.2 \times 10^6$ | |
| LwaCas13a-2 | RNA | ssRNA | 2 | 1800 | $1.1 \times 10^6$ | |
| LbuCas13a-1 | RNA | F-UUUUU-Q | 23 | 5800 | $4.0 \times 10^6$ | |
| LbuCas13a-2 | RNA | | 11 | 5600 | $2.0 \times 10^6$ | |
| SuCas12a2 | | ssRNA<br>F-UUUUU-Q | 0.124 | 256 | $4.9 \times 10^5$ | |
| SuCas12a2 | RNA | ssDNA<br>F-TTATT-Q | 0.113 | 131 | $8.6 \times 10^5$ | **Jang et al, 2025** |
| SuCas12a2 | | dsDNA<br>F-AGAACCGAATTTGTGTAGCTTATCAGACTG<br>Q-TCTTGGCTTAAACACATCGAATAGTCTGAC | 0.001 | 61 | $1.6 \times 10^4$ | |
| SuCas12a2 | | ssRNA<br>F-UGUAACGACC-Q | 0.026 | 90 | $2.9 \times 10^5$ | |
| SuCas12a2 | | ssDNA<br>F-TGTAACGACC-Q | 0.0123 | 25 | $4.9 \times 10^5$ | |
| SuCas12a2 | RNA | dsDNA<br>F-TGTAACGACC-Q<br>ACATTGCTGG | 0.012 | 42 | $2.9 \times 10^5$ | **This work** |
| SuCas12a2 | | dsDNA<br>F-TGTAACGACC<br>ACATTGCTGG -Q | 0.0145 | 40 | $3.6 \times 10^5$ | |

| Supplemental Table 5. Raw endpoint data for Cas12a2 and Cas13a activity plot in presence of total yeast RNA | | | | | | | |
|---|---|---|---|---|---|---|---|
| Total yeast RNA (ng/ul) | Target RNA (nM) | Fluorescence Intensities (RFU) | | | | | |
| | | Cas12a2 (Replicates) | | | Cas13a (Replicates) | | |
| 0 | 50 | 2260 | 2413 | 2425 | 51 | 61 | 68 |
| 1000 | 0 | 77 | 39 | 45 | 5 | 0 | 10 |
| 1000 | 50 | 2081 | 2333 | 2517 | 0 | 8 | 0 |
| 100 | 50 | 2319 | 2348 | 2287 | 19 | 0 | 0 |
| 10 | 50 | 2385 | 2416 | 2545 | 34 | 31 | 30 |
| 1 | 50 | 2574 | 2592 | 2395 | 52 | 53 | 53 |
| 0.1 | 50 | 2369 | 2358 | 2474 | 50 | 59 | 47 |
